# The origin of secondary structure transitions and peptide self-assembly propensity in trifluoroethanol-water mixtures

**DOI:** 10.1101/2023.09.14.557831

**Authors:** Anup Kumar Prasad, Rajarshi Samajdar, Ajay Singh Panwar, Lisandra L. Martin

**Author notes:** This work was conducted while this author was doing an internship at Indian Institute of Technology Bombay (Rajarshi Samajdar).

## Abstract

The formation of transient helical intermediates, implicated in the early-stages of amyloid formation in amyloidogenic peptides, is thought to be enhanced by membrane-peptide interactions. Uperin 3.5 is a seventeen-residue antimicrobial, amyloidogenic peptide that forms amyloid in phosphate buffered saline (PBS). The role of 2,2,2-trifluoroethanol (TFE) concentration, a known α-helical stabiliser, in modulating aggregation of Uperin 3.5 peptide in membrane-mimetic TFE:water mixtures was investigated. Thioflavin T (ThT) fluorescence assays showed complete inhibition of aggregation at higher concentrations of TFE (≥ 20% TFE:water v/v). However, a five-to-seven-fold increase in fibrillation kinetics was observed at 10% TFE:water mixtures in comparison to aggregation in a buffer. Further, aggregation in TFE:water mixtures was only observed upon addition of buffer. Interestingly, circular dichroism (CD) spectra showed the appearance of partial helical structures in 10% TFE:water, which transitioned to β-sheet rich structures only after addition of buffer. Microsecond time-scale molecular dynamics (MD) simulations of multiple U3.5 peptides in both salt-free and salt-containing TFE:water mixtures showed that changes in the local environment of peptide residues determined the structural transition and aggregation trajectories for U3.5. Consistent with experiments, the greatest extent of aggregation was observed for low TFE concentration (10% TFE:water simulations), characterised by faster formation of helical intermediates (oligomers). While the presence of 10% TFE efficiently induced partial helical structure in individual U3.5 peptides, it did not impede peptide-peptide interactions, thus enabling peptide aggregation. Addition of salt, screened like-charge repulsion between positively charged residues of different peptides, leading to stronger inter-peptide interactions. Significantly, the presence of salt determined subsequent structural transitions in the helical intermediates; either forming a predominantly α-helical oligomer in salt-free solutions or a β-sheet-rich oligomer in salt-containing solutions.

## Introduction

Many proteins can self-assemble into highly ordered, beta-sheet-rich fibrils, known as amyloids. Amyloid formation is associated with several human diseases, including Alzheimer’s disease, Parkinson’s disease and Type-2 diabetes.^1,2^ The generally accepted model used to describe fibrillation kinetics is the nucleation-elongation mechanism. In this model, the fibrillation kinetics pathway is characterised by an initial lag phase which involves nucleation of oligomeric species composed of partially folded or misfolded proteins.^3,4^ The lag phase is followed by an elongation phase involving rapid growth of fibrils, formed as a result of monomer addition to stable nuclei through the process of self-assembly. Finally, fibrillation kinetics reaches a saturation phase when equilibrium is achieved between the respective association and dissociation rates of monomer addition to the growing fibrils.^5^ Furthermore, these intermediate oligomeric species of amyloid-forming proteins, have been shown to be more cytotoxic than their mature fibrils.^6,7^ The degree of toxicity also varies among different oligomers of the same protein, possibly due to differences in structural and morphological features.^8–11^ For instance, oligomer toxicity may depend on their hydrophobic surface area and overall size which could determine membrane-oligomer interactions and diffusion of oligomers across a membrane.^12^ Oligomeric species affect cellular toxicity of amyloids and also determine the kinetics of fibril formation. Thus, the oligomerisation process continues to attract considerable attention with an aim to develop potential therapeutics for amyloidosis. 2,2,2-trifluoroethanol (TFE) is one of the most common cosolvents used to study conformational transitions of proteins and peptides.^13^ The ability of TFE to induce secondary structure, primarily the alpha-helix, makes it suitable for conformational studies of disordered proteins, as well as globular proteins.^13–15^ The structural transitions in proteins have been reported as a result of several factors such as, an increased intramolecular interaction in proteins^16^, lowering of the dielectric constant in an aqueous solution^17^, preferential clustering of TFE around a protein as well as entropic and enthalpic contributions.^18^ TFE molecule exhibits amphiphilic nature, which is made up of the hydrophilic hydroxyl group (OH) and the hydrophobic trifluromethyl (CF_3_) group. Depending on environmental factors, such as concentration and temperature, TFE can either interact directly with a protein or it can form small micelle-like clusters by self-association which can surround a protein.^19^ TFE addition into aqueous solution increases hydrophobicity and decreases the dielectric constant of the mixture. This can change hydrophobic and electrostatic interactions for a protein in a TFE-water mixture, thus significantly alter protein folding and aggregation. Water-TFE mixtures can provide a lipophilic and lipophobic interface for a protein which mimics membrane-water interfaces.^20^ Thus, TFE has been an attractive option for protein and peptide folding studies and provides certain advantages over lipid-based membranes. For instance, TFE can overcome a line broadening effect that is induced by peptide-micelles in NMR.^13,20,21^

Uperin 3.5 (U3.5) peptide is an antimicrobial peptide (AMP), secreted onto the skin of Australian toadlet (*Uperoleia mjobergii*), which can self-assemble into amyloid in buffered saline.^22,23^ U3.5 displays a variety of secondary structure preferences depending on solvent type and solvent conditions; such as random coil in pure water, alpha-helical in bacterial mimetic environments, and beta-sheets in saline buffer.^24–26^ Seventeen residues long, the U3.5 peptide is cationic in nature with amidation at the C-termini and can form an amphipathic α-helix. The overall charge and amphipathic properties of U3.5 are important in determining peptide-peptide and peptide-lipid interactions, and in determining the peptide’s functional role as an antimicrobial peptide. Amyloid-β, α-synuclein and many other amyloid forming peptides have shown toxic oligomerisation at peptide-lipid interface. The transitions from soluble toxic oligomers to amyloidogenic nuclei and vice versa are key processes that exhibit two important characteristics: firstly peptide stability by amyloid formation in absence of membrane and secondly cellular toxicity by toxic oligomers in presence of membrane. Understanding the molecular basis of the structural transitions, enables discovery of specific targets needed for inhibition of peptide oligomerisation (or self-assembly) into pathogenic disease states. The ability of U3.5 peptide to acquire various secondary structures in different solvent conditions makes it an ideal candidate to investigate structural transitions in soluble oligomers and their role in amyloid formations.

The current study combined experimental and MD simulation techniques to investigate the mechanisms of peptide oligomerisation of U3.5 peptide in TFE-water mixtures, and its influence on amyloid formation. Whereas, amyloid formation of U3.5 peptide significantly increased with the addition of smaller amounts of TFE in water, U3.5 aggregation was completely inhibited at higher TFE concentrations.^25^ We employed TFE:water (v/v) ratios of 10%, 15%, 20% and 40% in presence and absence of PBS. The spectroscopic methods of Circular dichroism (CD) and Fluorescence assay using Thioflavin T (ThT) were used to assess time-dependent secondary structure and amyloid formation kinetics of U3.5 peptide, respectively. MD simulations revealed the molecular basis of peptide oligomerisation and its role in β-sheet-rich aggregation.

## Materials and methods

### Experimental method

#### Materials

U3.5 peptide, amidated at the C-terminus was purchased from Peptide 2.0 Inc. (Chantilly, VA). Potassium phosphate monobasic (KH_2_PO_4_, anhydrous 99%) and potassium phosphate dibasic (K_2_HPO_4_, anhydrous), purchased from Sigma-Aldrich (St. Louis, USA), were used in the preparation of phosphate-buffered saline (PBS) solution. Sodium chloride (NaCl ≥ 99.5%) was also used in the PBS solution. Ultrapure water (18.2 MΩ cm) was used for buffer preparation, peptide dissolution, etc in all experiments. The pH of the buffer was adjusted to 7.4 ± 0.05 by addition of sodium hydroxide (NaOH). Using these materials, a 10-fold concentrated PBS solution (10X) was prepared and filtered using polypropylene membrane filters (GH Polypro, 0.2 μm, PALL Life Sciences Corp., Port Washington, NY). In all the experiments, 10X PBS was used to make a final concentration of 20 mM phosphate and 100 mM NaCl. The thioflavin T (ThT, Sigma-Aldrich) was diluted in dimethyl sulfoxide (DMSO) to make 1 mM stock solution. The ThT stock was protected from light and stored at -20 °C. The final concentration of ThT in assay was 10 µM.

#### Fluorescence (Thioflavin T) Assays

U3.5 peptide has shown a variety of secondary structures and aggregation status in different solvents. The U3.5 peptide acquires random coiled structure in water so, it was diluted in ultrapure water to prepare 500 mM stock solutions. These stock solutions were stored at -20 °C prior to their use in experiments. TFE was diluted using ultrapure water to make a 50% TFE:water (v/v) stock solution, and added to the samples in order to gain the required final percentage TFE:water (v/v). The solvent components were added to the samples in chronological order required to study the secondary structure transitions and aggregation with minimum perturbation of the peptide samples. The peptide concentration varied a little due to the addition of solvent components at different stages of experiments. ThT stock (1 mM) was prepared in water, which was stored at −20 °C with protection from the light. The final concentration of ThT in each sample was 20 mM. 96-well microplates (black polystyrene) with bottom optic, non-binding surfaces from Greiner Bio-One (Germany) were used for the fluorescence ThT kinetic assay. Fluorescence was measured using a CLARIOstar plate reader (BMG Labtech, Germany) with excitation and emission wavelengths of 440 and 480 nm, respectively. All experiments were carried out with 5 min measurement gap at 37°C under quiescent conditions. These experiments used a final volume 100 µl and were performed in triplicate.

#### Circular Dichroism

Samples were prepared, combined in chronological order using the same protocol described for the Fluorescence ThT assay. J-815 Circular dichroism (CD) spectropolarimeter (Jasco Corp., Tokyo, Japan) and a quartz cuvette (21/10/Q/1/ CD, Starna Scientific Ltd., Essex, UK) of 1 mm path length were used for CD experiments. Far-UV CD spectra (195−260 nm) were recorded at standard sensitivity (DIT, 1 s; bandwidth, 1 nm; data pitch, 0.5 nm; continuous; scanning speed, 50 nm/min; single scan). CD measurements were performed for 12 hrs to 24 hrs after addition of each solvent component, as required. Spectra were recorded every minute for the first one hour following addition of a component and thereafter every 30 minutes until the next solvent component was added. CD spectra for buffer and buffer + TFE solutions were also recorded for baseline correction of each sample spectrum.

#### Molecular dynamics simulations

Fully-atomistic MD simulations were carried out using the CHARMM36 force field with the NAMD simulation package. Simulation systems were modelled to study U3.5 peptide in TFE:water mixtures in both salt and salt-free conditions, respectively. Two categories of MD simulations were setup; (i) monomer simulations, involving single U3.5 peptides in TFE:water mixtures, and (ii) multi-peptide simulations, involving four U3.5 peptides in TFE:water mixtures. The primary objective of the monomer simulations was to understand the effect of TFE composition on peptide secondary structure. Multi-peptide simulations were designed to understand the effect of TFE composition on peptide aggregation and secondary structure transitions during aggregation. Since the C-terminus of U3.5 is naturally amidated, U3.5 peptides in the simulations were also modelled with an amidated C-termini.

U3.5 peptide is randomly placed in a cubic simulation box (of 60 Å^3^ for monomer and 100 Å^3^ for multi-peptide simulations) in TFE:water mixtures with varying TFE concentrations without salt and with salt (concentration = 0.15 mM) present. Additional details of simulations are provided in Table S1. In all the cases, a random coiled conformation of the U3.5 peptide (generated with VMD) was used to setup the simulations systems. Periodic boundary conditions were applied along all three directions of the simulation box to model a bulk system. Electrostatic interactions were calculated using Particle-mesh Ewald method with a grid spacing of 1 Å. A switching function was used for smooth decay of Lennard-Jones interaction in range of 10 Å to 12 Å. Temperature and pressure in the simulations were maintained by a Langevin thermostat and a Nosé-Hoover Langevin piston, respectively. All systems were simulated under physiological conditions with 310 K temperature and a 2 fs timestep. Relaxations of simulations were done by applying harmonic constraints on peptides so that solvent molecules could relaxed around the peptides. The harmonic constraints were gradually removed in 2 ns and systems were further minimised for 400 ps of simulation without any constraints. Systems were initially equilibrated in an NVT ensemble (1 ns, *T* = 310 K), followed by equilibration in an NPT ensemble (1 ns, *T* = 310 K, P = 1 atm). All production runs were conducted in an NVT ensemble at 310 K.

## Results and Discussion

### Experimental Approaches

#### Peptide aggregation kinetics

Our previous studies on U3.5 peptide revealed amyloid fibril formation in saline buffer, which disintegrated on addition of sodium dodecyl sulphate (SDS).^27^ Various membrane mimetics (TFE, SDS, or liposome of POPC) have been reported to induce helical peptide secondary structures and influence aggregation kinetics of U3.5 peptide (typically observed in saline buffer).^24,27,28^ The aggregation kinetics of an amyloidogenic peptide, such as U3.5, in a membrane mimetic environment will be influenced by multiple factors, including peptide-solvent interactions, solvent concentration and presence of electrolyte in the medium. Previous studies of U3.5 aggregation in presence of membrane mimetics have not attempted to decouple the individual effects of these factors. In addition, U3.5 (and other uperins) tend to adopt helical conformations in contact with membrane mimetic or more hydrophobic environments, which typically suppresses aggregation and amyloid formation. In this context, the current study raises the following pertinent questions. Firstly, how does TFE, a membrane mimetic and α-helix stabiliser, alter U3.5 aggregation kinetics both in presence and absence of saline buffer? Secondly, what are the individual roles of TFE and salt in modifying U3.5 aggregation kinetics in membrane mimetic TFE:water mixtures? Thirdly, what are the molecular mechanisms responsible for modifying U3.5 aggregation kinetics in TFE:water mixtures?

In the current study, sequential addition of TFE and saline buffer to a U3.5 sample was used to assess their respective, individual effects on U3.5 aggregation kinetics and secondary structure evolution.

Figure 1 shows an aggregation (kinetic) profiles of U3.5 peptide at various TFE volume percentages in aqueous solution. Two different fluorescence ThT assays were carried out with different purposes. In the first assay, TFE and buffer were sequentially added to the sample to investigate their individual effects on aggregation (Figure 1(a)). In the second assay, TFE and buffer were added together to the sample to assess whether aggregation kinetics differed from the first assay (Figure 1(b)). U3.5 samples, initially prepared in water (111 μM), did not show any aggregation. The peptide remained in an unaggregated state even after addition of TFE at 4 hours (Figure 1(a)). However, the profile changed dramatically after buffer was added at 23 hours. As anticipated, the peptide started to self-assemble immediately following addition of saline buffer into the samples. Surprisingly, the TFE content significantly impacted the aggregation kinetics of U3.5. The lower amounts of TFE (5% and 10%) showed a five- and seven-fold increase in saturation intensities, respectively, when compared to the 0% TFE case. In contrast, solutions containing higher TFE percentages amyloid formation was completely inhibited in U3.5 peptide, although 20% TFE lay a few percent above the zero value. In the second ThT assay, TFE:water mixtures with varying TFE concentrations were prepared in saline buffer and then added to U3.5 peptide samples. In this case, the samples showed the same aggregation kinetics as obtained in first ThT assay after addition of buffer (Figure 1(b)). In the second set of assays, an excess of TFE was added to the 5%, 10% and 20% TFE samples, to increase TFE concentration to 40% v/v, at 23 hours. In each case, this resulted in a complete loss of fluorescence intensity in all samples, indicating disintegration of amyloid fibrils in excess TFE. In both experiments, 10% TFE-water increased aggregation seven-fold for U3.5 compared to the peptide sample in buffer alone. These data indicated a significant effect of the TFE content on peptide aggregation; although importantly, there was no ThT fluorescence in media containing TFE until buffer was added to these samples. Hence, a structural analysis of U3.5 peptide was undertaken to investigate the influence of TFE solvent at different stages of U3.5 peptide aggregation.

**Figure 1.**
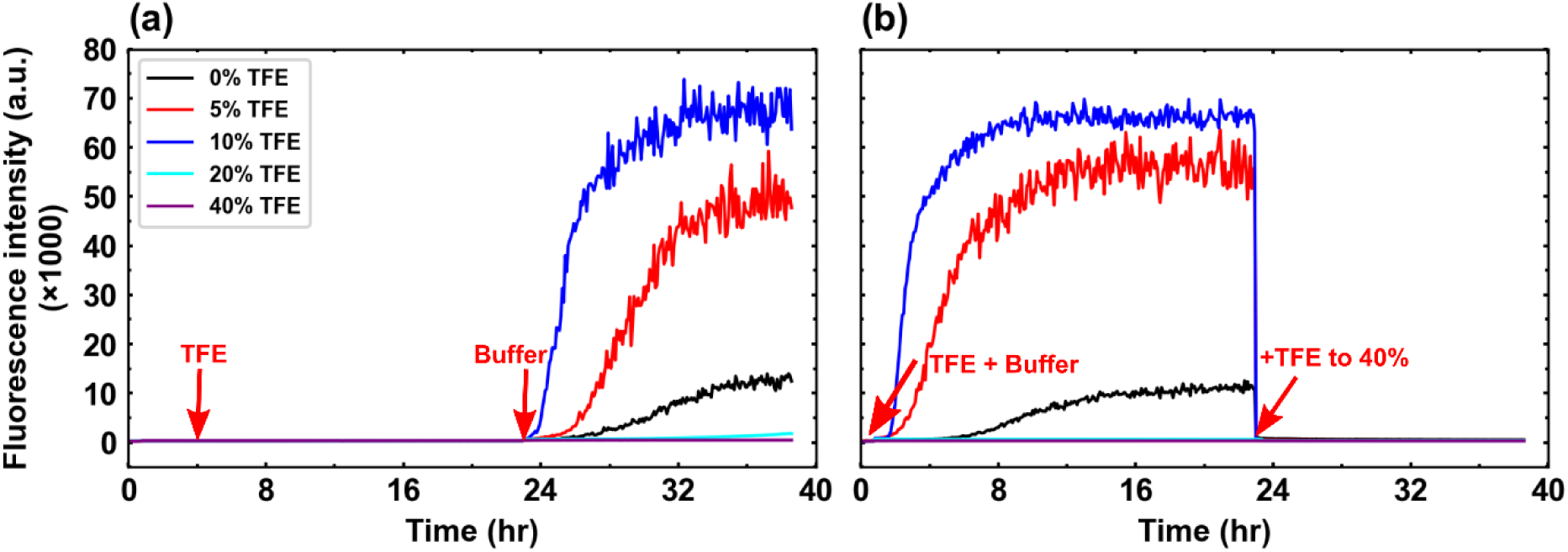
ThT fluorescence showing aggregation kinetics of U3.5 with sequential addition of TFE and buffer. (a) Peptide samples of 100 µM concentration, prepared in water, did not show any aggregation even after addition of TFE at 4 hrs of the experiment. However, addition of PBS buffer at 23 hours stimulated peptide aggregation with a significant variation in kinetics with respect to the TFE concentration. (b) TFE and buffer added together to the sample resulted in aggregation profiles similar to those in (a) after addition of buffer. Addition of an excess amount of TFE (up to 40%) at 23 hours led complete loss of fluorescence in all sample, which indicated the disintegration of amyloid.

#### Secondary structure transitions

Many proteins, such as Aβ, α-synuclein, transthyretin and lysozyme, form highly ordered amyloid fibrils that are unrelated in their native secondary structure.^29,30^ The phenomena of structural transition of proteins from their native secondary structures to β-sheet-rich amyloid fibrils suggests a correlation between structural transitions and amyloid formation. The variation in aggregation kinetics at different TFE concentrations (Figure 1) indicated that secondary structure transitions in TFE may be crucial to understanding aggregation kinetics. CD measurements were carried out at specific time intervals to capture the structural evolution of the U3.5 at different stages of peptide aggregation. U3.5 showed a random coil structure in water (black curves, Figure 2(a)), which transitioned into partial helical structures after addition of 10% TFE (red curves, Figure 2(a)). This implied that a solution containing a small amount of TFE could induce partial helical structures of the U3.5, although did not result in fibril formation. Interestingly, addition of saline buffer to this mixture resulted in the transition of the partial helical secondary structure into β-sheet structures (Figure 2(b)). Deposition of amyloid fibrils on the wall of the CD cuvette was quite high compared to fibril formation under buffer-only conditions (Figure S1). Further addition of TFE to a final concentration of 40% caused disintegration of U3.5 amyloid fibrils, which was accompanied with structural transitions from β-sheet to highly helical structures (Figure 2(c) and Figure S1). In a separate experiment, 40% TFE was added to a U3.5 sample in water (Figure 2(d-f)). The resulting spectra showed a rapid transition from random coil to helical structure (Figure 2(e)). These spectra were similar to those observed in the spectra obtained in Figure 2(c). The spectra showed unchanged secondary structure of peptide on addition of saline buffer into this sample (Figure 2(f)).

**Figure 2.**
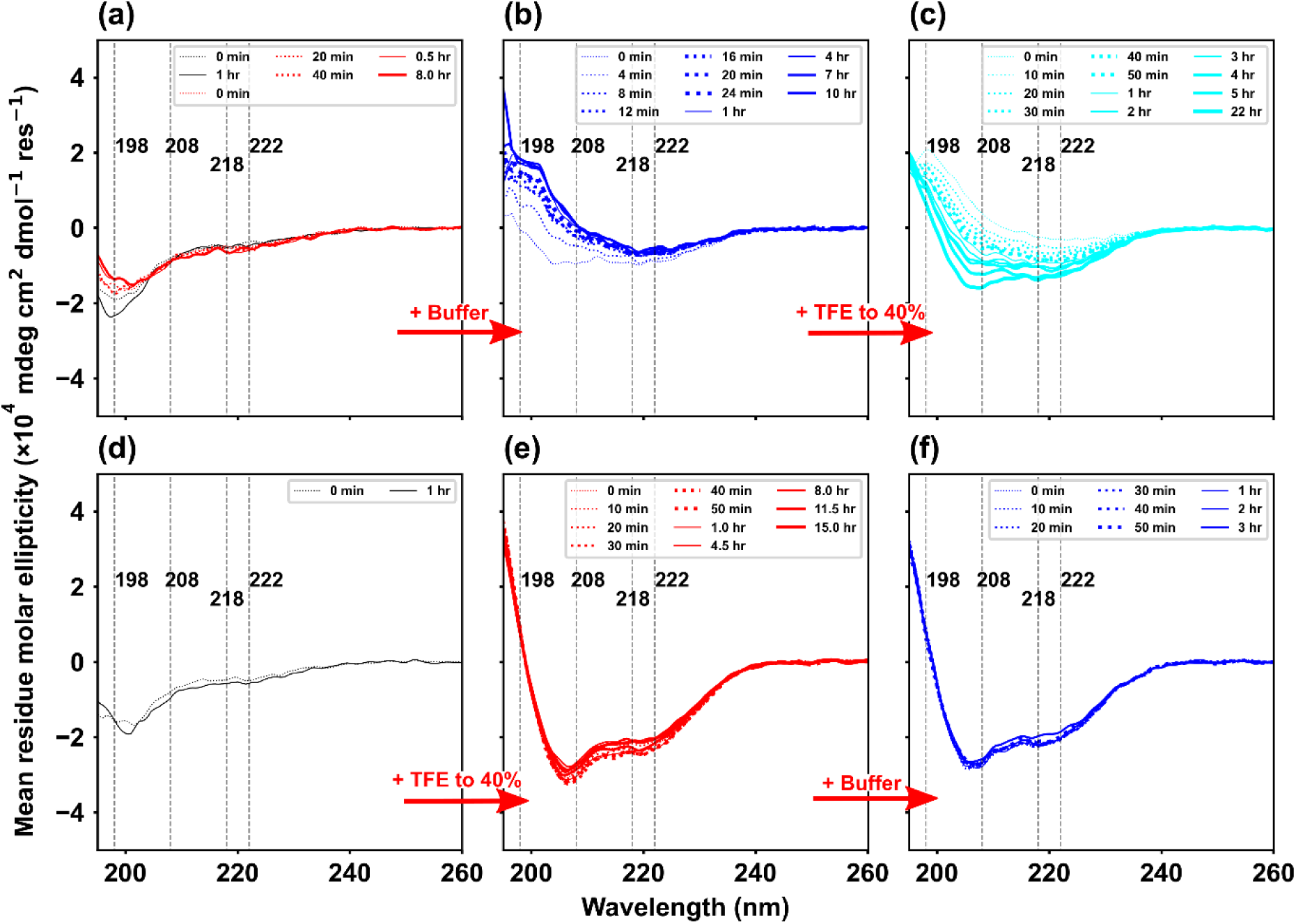
Time-dependent CD spectra of U3.5 peptide (100 µM) on sequential addition of TFE and PBS buffer. The CD spectra before and after of an hour are shown in dotted and solid curves, respectively, with an increasing curve thickness corresponding to the time points. (a) The CD spectra (black) indicated that the U3.5 was initially in a random coil structure, which transitioned into partial helical structures (red) on addition of 10% TFE (v/v). (b) U3.5 peptide quicky transitioned into β-sheet secondary structures, within 10 minutes, on addition of PBS buffer to the samples. (c) Further addition of TFE (to 40% v/v) resulted β-sheet to α-helix transition in U3.5 peptide. (d) Another experiment of U3.5 peptide with random coil structure (black, in water) showed rapid transition on addition of 40% TFE (panel d → e, compared with panels b → c), which were not affected by addition of PBS buffer (f).

The transient appearance of helical structures, prior to formation of β-sheet-rich amyloid fibrils, has been previously reported for several proteins, including Aβ peptide, α-synuclein, and islet amyloid polypeptide (IAPP, amylin).^24,31–34^ However, to date, the connection between secondary structures and aggregation kinetics is not well understood. Figures 1 and 2 provide a clear correlation between secondary structure transitions and different stages of peptide aggregation. In Figure 1, the aggregation of U3.5 in buffer alone showed a longer lag phase (≈ 2 – 4 hrs) compared to the kinetics at 5% TFE (lag phase ≈ 30 minutes – 1 hr) and 10% TFE (no lag phase), respectively. Secondary structure transitions of U3.5 in buffer (Figure S2) indicated that slow amyloid formation kinetics was accompanied by a slow transition from random coil to β-sheets (Figure 1 and Figure S2). In contrast, the aggregation kinetics of U3.5 at 5% and 10% TFE started the elongation phase almost immediately upon addition of saline buffer to the sample (Figure 1). The corresponding secondary structure transitions at 10% TFE can been seen in CD spectra of Figure 2(a) and (b). The partial helical structure of peptide (red curve), induced by 10% TFE (Figure 2(a)), rapidly transitioned into a β-sheet secondary structure, within 10 minutes of buffer addition (Figure 2(b)). The aggregation kinetics and corresponding structural transitions of U3.5 implied that a reduced lag phase and increased aggregation are associated with partial helical structures induced at low TFE concentrations. Prefibrillar and soluble oligomers, are thought to be formed in the lag phase of peptide aggregation and play a crucial role in aggregation kinetics of amyloid.^35^ The combined results obtained from ThT assays and CD spectra suggest that the low concentration of TFE may have induced formation of peptide oligomers (containing partial helical structure) prior to addition of buffer. This would have resulted in the shortened lag phase observed at low TFE concentrations (5% and 10% TFE). We have previously observed these partial helical structures in MD simulations of U3.5 aggregation in salt-containing aqueous solutions, which highlighted the role of helical intermediates in β-sheet-rich aggregation.^36,37^ Thus, to better understand the molecular mechanism at the early stages of aggregation; MD simulations were carried out using different TFE concentrations in salt-free and salt solutions.

## Molecular dynamics simulations

### Monomer simulations

A single U3.5 peptide was simulated in TFE:water mixtures with varying TFE compositions (0 – 40% TFE), to understand the effect of TFE composition on structural transitions at the level of an isolated peptide (Figure 3(a)). Simulations were carried out in both salt-free and salt (0.15 M NaCl) solutions. The initial states of these simulations corresponded to single U3.5 peptides in random coil conformations. Changes in various secondary structure components (with a focus on α-helix and β-sheet) were tracked over the course of the simulation trajectories. Figure 3(a) shows snapshots of induced helical structure of U3.5 peptide simulated at various TFE:water systems. It also highlights the change in TFE distribution in the immediate vicinity of the U3.5 peptide at different TFE concentrations. An increase in TFE concentration was associated with increased helical content in the peptide secondary and increased crowding of TFE around the peptide. The increase in α-helical structure with increasing TFE content was observed for both salt-free and salt-containing cases. This trend is plotted in Figure 3(b) and is consistent with the role of TFE as a known α-helix stabiliser.

**Figure 3.**
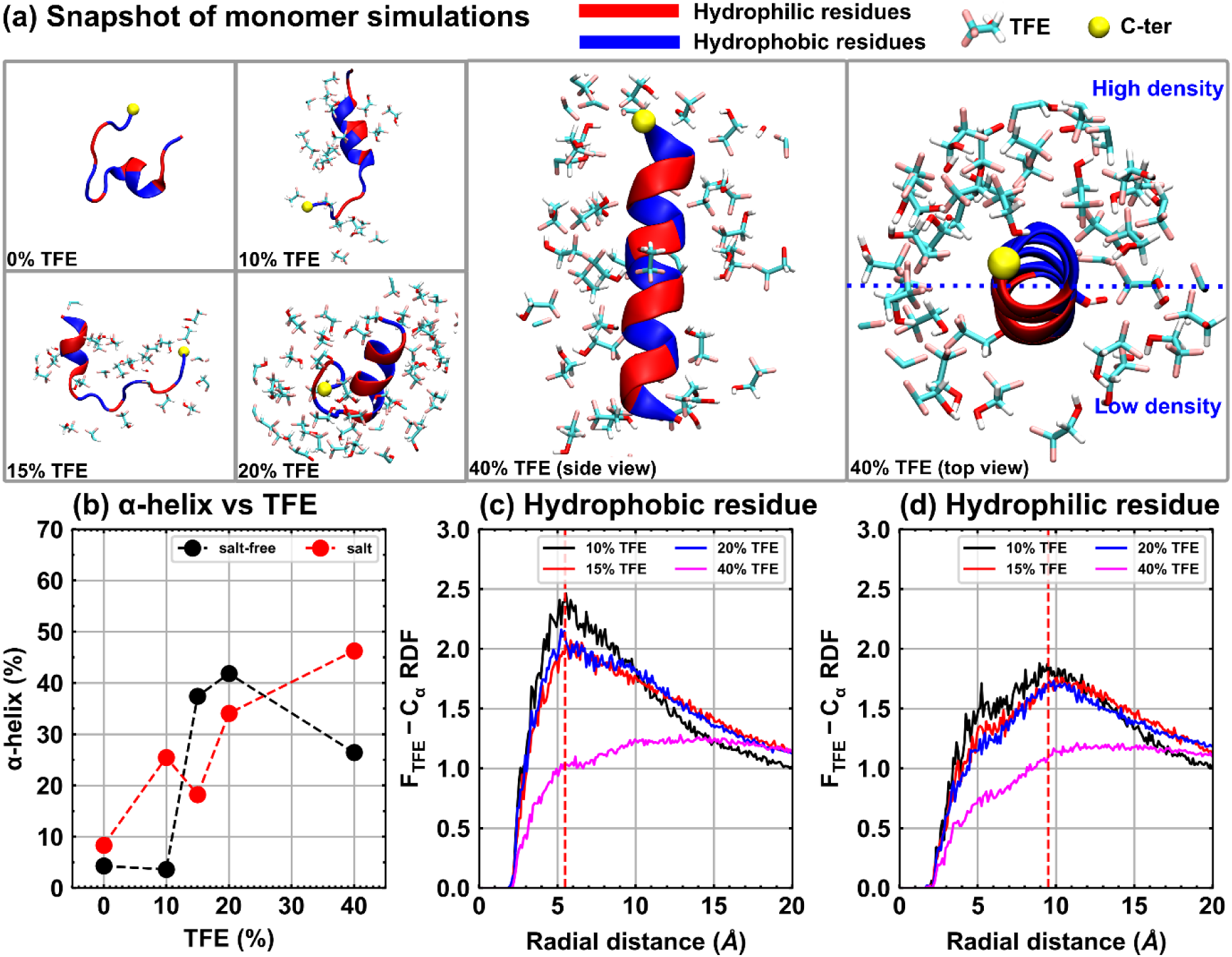
Evolution of U3.5 secondary structure and TFE distribution around the peptide. (a) Snapshots of induced helical structure and TFE distribution around U3.5. TFE is strongly associated with hydrophobic residues (blue) compared to the polar residues (red). (b) Correlation between α-helix of U3.5 peptide and TFE percentage in the aqueous solution, plotted based on averaged α-helices in the last 200 ns of triplicate trajectories. The helical content in the peptide were increased in both salt-free and salt cases of monomer simulations. Radial distribution functions (RDF) of TFE (fluorine atom) were calculated around alpha carbon of hydrophobic residues (c) and hydrophilic residues (d). TFE molecules are strongly associated with hydrophobic residues with solvation shells of radius 5.5 Å, whereas the TFE molecules are pushed by water molecules at 9.5 Å around hydrophilic residues.

The evolution of α-helical secondary structure in the individual simulation trajectories is shown in Figure S3. Visual inspection of simulation trajectories indicated preferential interactions of hydrophobic residues with TFE molecules and hydrophilic residues with water molecules, respectively. Peptide-TFE interactions were quantified by calculating radial distribution functions (RDF) corresponding to the distribution of fluorine atoms (F of TFE) around the alpha carbon of the peptide (C_α_). Figures 3(c) and 3(d) show RDF plots of TFE (fluorine) distribution with respect to C_α_, averaged over all hydrophobic and hydrophilic residues, respectively. The peak RDF values (corresponding to the first solvation shell at 5.5 Å) around the hydrophobic residues were 2 – 2.5 times greater than the bulk concentration (Figure 3(c)). In contrast, RDF values in the immediate vicinity of the hydrophilic residues were lower and the first TFE peak was shifted to 9.5 Å away from the C_α_ atoms (Figure 3(d)). However, RDF plots corresponding to 40% TFE deviated from the above behaviour. Importantly, systems with 40% TFE were characterised by a homogeneous distribution of TFE molecules around both hydrophobic and hydrophilic residues due to the increased fraction within the solution. Thus, the monomer simulations provided insight into the interactions of TFE with the peptide and their impact on the resulting peptide secondary structure. However, the effects of TFE on peptide aggregation and structural population evolution of aggregates required further investigation.

### Multi-peptide simulations

#### Secondary structure

Monomer simulations revealed secondary structure changes in a single U3.5 peptide with increasing TFE concentration in aqueous solutions. Multi-peptide systems with four U3.5 peptides in TFE:water mixtures, for cases of both salt-free and salt-containing solutions, were simulated to investigate U3.5 aggregation as a function of TFE concentration. The α-helix content at all concentrations of TFE increased with time over the course of the 500 ns trajectories. This was true for both salt-free and salt containing solutions (Figures 4(a) and 4(d)). Every trajectory shown here was averaged over three independent trajectories. The secondary structure transitions for the independent trajectories are shown in Figure S4. Interestingly, β-sheet transitions in the peptide systems appeared to depend on the presence of salt in the TFE:water mixtures. For salt-free solutions, a significant amount of β-sheet (≈10% of the secondary structure) was observed only for 0% TFE (pure water). The β-sheet content was negligible for all finite TFE concentrations considered in the simulations (Figure 4(b)). These results were consistent with the monomer simulations from Figure 3, where TFE primarily promoted transitions to α-helix dominated secondary structures in U3.5. The results also reinforced the conventional view of TFE as an α-helix stabiliser in peptides. The variations of both α-helix and β-sheet components are plotted as functions of TFE concentration in Figure 4(c). The values plotted in Figure 4(c) represent time averages taken over the last 200 ns of the trajectories in Figures 4(a) and (b). Although, the α-helix content increased with TFE concentration, it passed through a maximum (nearly 35% α-helix) at 10% TFE, before saturating to approximately 20% α-helix at higher concentrations. On the other hand, the β- sheet content rapidly disappeared upon addition of TFE in the mixture.

**Figure 4.**
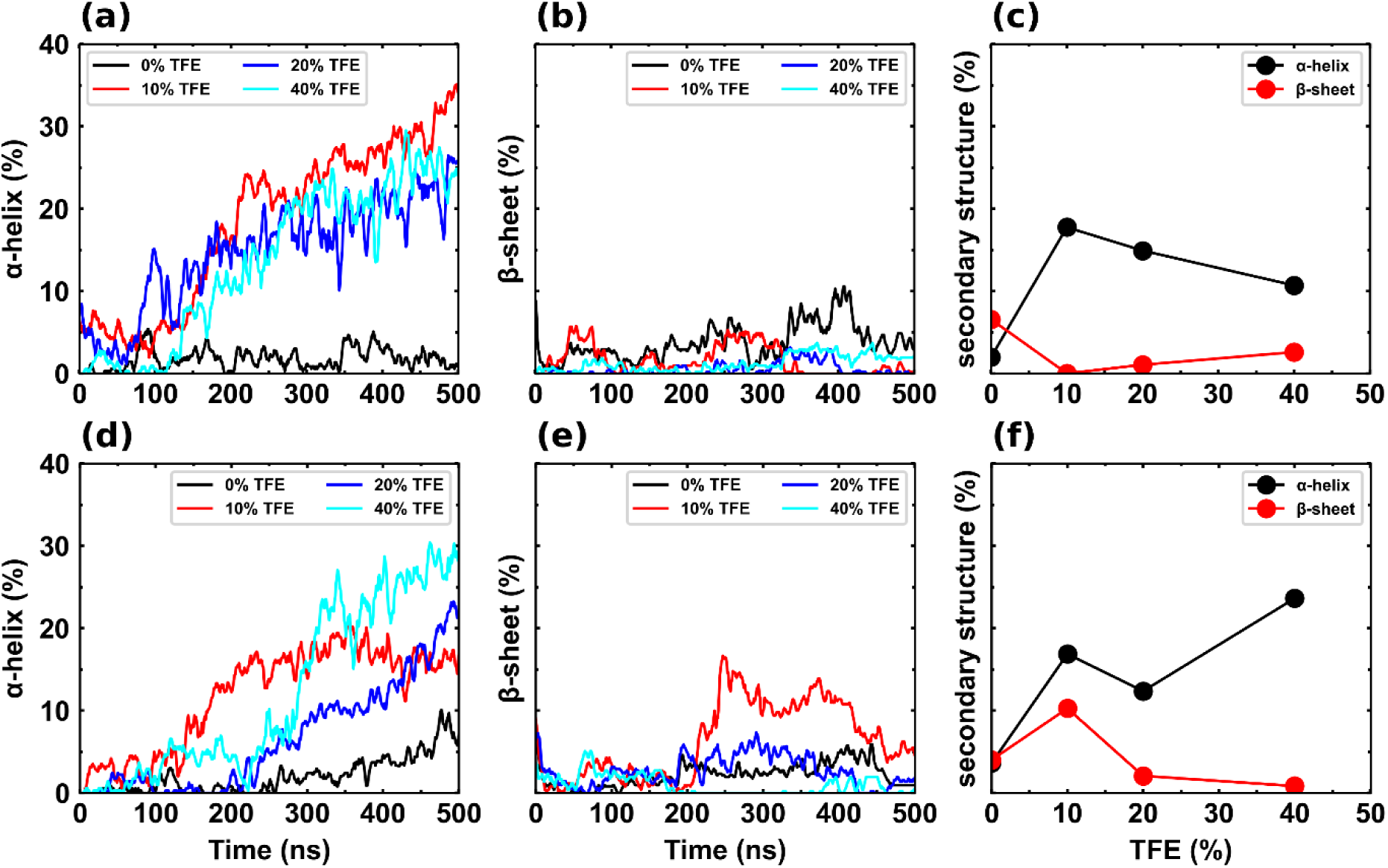
Secondary structure evolution in U3.5 peptide and correlations between TFE concentrations and induced structural contents of muti-peptide simulations in salt-free and solutions. Each data curve is an average of three replicates of corresponding simulation systems. The correlation between TFE % and secondary structure were plotted based on the last 300 ns of the trajectories. In salt-free systems, the helical content of peptides increased with the incremental percentage of TFE (a), whereas the β-sheet content was negligible in all TFE:water simulations (b). Similarly, in salt containing muti-peptide simulations, the U3.5 peptide showed the increasing trend in helical content with increasing TFE % (d). However, the β-sheet formation was only significant (formed up to 15%) in U3.5 of 10% TFE (e). The correlation of secondary structure transitions of U3.5 with TFE percentage are shown for salt-free (c) and salt (f) solutions.

However, a qualitative change was observed upon addition of salt to the TFE:water systems. Similar to the salt-free solutions, the α-helix amount increased with increasing TFE concentration. However, the prominent maximum in α-helix amount observed in Figure 4(c) was not seen for salt solutions (Figure 4(f)). Instead, α-helix amounts seemed to be nearly 20% for all TFE concentrations (within error). The trajectories in Figure 4(e) indicated higher amounts of β-sheet content when compared to the salt-free case. Indeed, Figure 4(f) clearly showed higher β-sheet content when compared to Figure 4(c). The most interesting observation was that the β-sheet content varied non-monotonically with increasing TFE concentrations, showing a maximum at an intermediate TFE concentration of 10%. This observation was in agreement with the experimental results in Figure 1, where the maximum ThT fluorescence intensity was observed at an intermediate TFE concentration of 10%. ThT fluorescence assays probe β-sheet containing amyloid structures^38^ in solution. Hence, the ThT results (Figure 1) showed that intermediate TFE concentrations (5 – 10%) resulted in rapid formation of β-sheet-rich self-assembled structures in solutions. The multi-peptide simulations in salt also showed a maximum in β-sheet content at 10% TFE suggesting that the presence of salt qualitatively altered secondary structure transitions in the peptides. In contrast with the single peptide simulations, inter-peptide interactions also affected secondary structure transitions when multiple peptides were present. Secondary structure transitions are expected to be determined not only by a change in the ionic environment of individual peptides, although these are likely to be influenced by peptide-peptide interactions. Further, peptide-peptide interactions are also modulated (screened) by the presence of salt in the solution. Secondary structure transitions in the multi-peptide systems are coupled with the process of peptide self-assembly. Hence, a detailed investigation of peptide aggregates (or clusters) formed in the multiple peptide simulations was needed to understand the molecular mechanisms leading to enhanced β-sheet content at intermediate TFE concentrations.

#### Peptide aggregation

Peptide aggregation in the multi-peptide simulations was quantified by counting the numbers of different sized clusters that formed over the course of a simulation. Since the simulation system comprised four peptides, the largest sized cluster was a tetramer. The numbers of dimers, trimers and tetramers that formed during a simulation are plotted with respect to time in Figure 5. Cluster formation in the simulation was identified by following a peptide-peptide distance criterion of 5 Å. Figure 5 clearly shows that cluster formation in U3.5 was strongly dependent on the TFE concentration. Initially, the overall cluster formation increased as the TFE concentration increased from 0 to 10% TFE. This was characterised by an increase in both frequency of cluster formation and the incidence of higher order clusters. Both sets of simulations (salt-free and salt solutions) for the 10% TFE case were extended to 1 µs to better characterize the peptide clusters. Interestingly, the multi-peptide simulations of 10% TFE showed highest peptide aggregation in both salt-free and salt solutions. An increase in TFE concentration beyond 10% led to reduced cluster formation for both the 20% and 40% TFE cases. In fact, the 40% TFE trajectories were predominantly characterised by U3.5 monomers, interspersed with transient occurrences of dimers. The individual cluster status over each trajectory for all data sets are shown in Figure S5.

**Figure 5.**
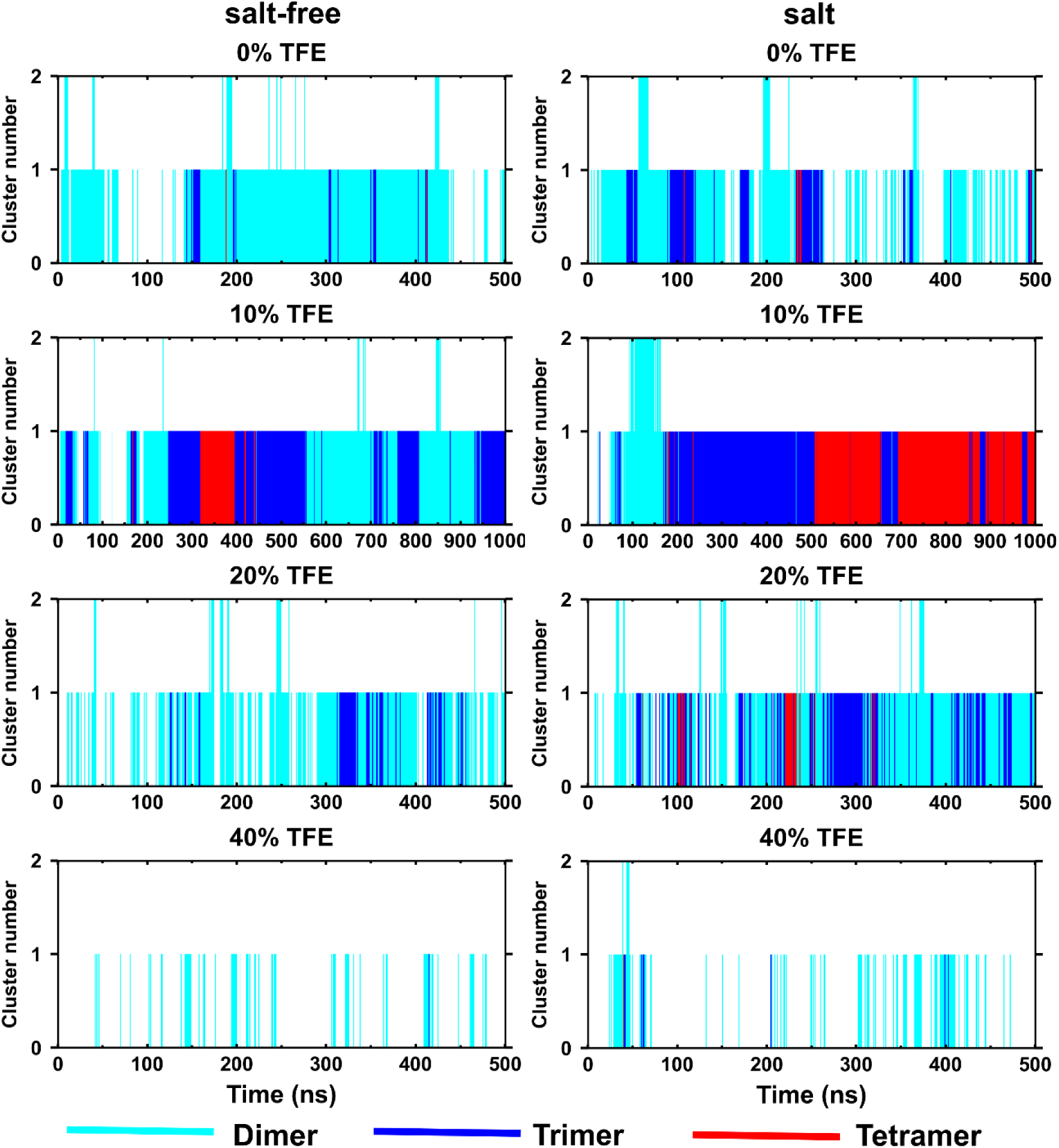
Heat map of U3.5 peptide clusters formed in multi-peptide simulations at varying TFE percentage. In both the cases, cluster formations increased at low TFE amount (10%) compared to no TFE (0%), which is further decreased in 20% TFE and then completely inhibited at 40% TFE.

In general, a larger number of higher order clusters were formed in presence of salt. This was evident for all TFE concentrations between 0 – 20% TFE. The effect of salt on peptide self-assembly was most evident when comparing the salt-free and salt solution clustering data for 10% TFE. TFE solutions with salt were characterised with many trimers and tetramers, implying a significant role of electrostatic screening in the self-assembly process. The plots also show steady growth of clusters by peptide aggregation in presence of salt. In presence of salt, at 10% TFE, two stable dimers were observed between 75 – 175 ns. One of these dimers grew into a trimer at approximately 175 ns, which existed as a stable trimer until approximately 500 ns. Further, the trimer grew into a stable tetramer after 500 ns. In contrast, dimers and trimers were less stable in the salt-free case, with the peptides switching between the dimeric and trimeric states.

The presence of salt in the 10% TFE mixtures resulted in the formation of more stable, higher order oligomers in comparison to the salt-free case. In addition, secondary structure evolution in the oligomers followed distinctly different trajectories for the salt-free and salt solutions, respectively (Figure 4). Whereas, the formation of oligomers was associated with a large increase in the α-helical content in the salt-free case (Figure 4(c)), oligomerisation was accompanied with significant increase in the β-sheet content only in presence of salt. During peptide oligomerisation, secondary structure transitions are coupled with inter-peptide interactions, and contribute to peptide self-assembly and fibrillation kinetics. The appearance of helical structures in the early stages of amyloid formation has been reported for many amyloid forming peptides, including U3.5.^24,31–33^ Our previous study on U3.5 peptide highlighted the role of helical intermediates (helix-rich oligomers) in the pathway to formation of β-sheet-rich aggregates. The observation of faster fibrillation kinetics (Figure 1) and enhanced β-sheet content (Figure 4(f)) at 10% TFE suggests that a certain amount of TFE may enable formation of helical intermediates on the pathway to formation of β-sheet-rich amyloids. It is important to understand the role TFE concentration (at the molecular level) in enabling formation of helical oligomeric intermediates and their subsequent transformation to β-sheet-rich aggregates. Hence, a detailed examination of secondary structure transitions, oligomerisation and inter-molecular interactions was undertaken for the special case of 10% TFE in both salt-free and salt solutions.

#### Evolution of helical intermediates

Oligomerisation is a crucial step in peptide aggregation, which produces small clusters of peptides (intermediates) in the lag phase of amyloid formation. Addition of seeds (preformed oligomers) to a solution of an amyloidogenic peptide (including U3.5) has been shown to decrease the lag phase, hence accelerate aggregation kinetics.^37,39^ Many amyloidogenic peptides have revealed diverse range of oligomers of different sizes and structures. The appearance of transient, helical structures in oligomers has been reported for many amyloidogenic peptides, including Aβ, α-synuclein and islet amyloid polypeptide (IAPP, amylin).^31,32,34,36,40,41^ In 10% TFE, the appearance of partial helical structures and faster aggregation kinetics, with a rather short lag phase, (Figure 2(a) and Figure 1, respectively) highlight the significance of helical intermediates on the pathway of amyloid formation. Both cases of multi-peptide simulations, salt-free and salt solutions, at 10% TFE concentration were extended to 1 µs. The helical components started increasing at around 200 ns for both sets of simulations. Oligomers with partial helical content (Figures S5 and 6) were observed in both cases. U3.5 peptides form amphipathic helices, and the oligomers were loosely bound clusters with hydrophobic cores. Structurally, these oligomers evolved further into either helix-rich or β-sheet-rich oligomers depending on absence and presence of salt in the solution, respectively (Figure 6). The compactness of peptide clusters was tracked by their radius of gyration, *R_g_* (Figure S6). Indeed, structural transitions from α-helix to β-sheet in salt solutions were accompanied with further compaction of U3.5 oligomers. Our previous study on U3.5 aggregation, in presence of salt, revealed that helical intermediates formed due to aggregation of amphipathic, U3.5 helices along their hydrophobic faces.^36^ The formation of a helical intermediate facilitated peptide-peptide interactions resulting in further structural transitions to β-sheet-rich U3.5 clusters. The helical intermediates, formed in 10% TFE simulations, evolved into two different oligomers depending on the presence or absence of salt in the solution. Whereas, α-helix-rich oligomers were observed in salt-free solutions, β-sheet-rich oligomers formed from the initial partial helical intermediates in presence of salt (Figure 6). These results indicated that helical intermediates formed as a first step toward aggregation and facilitated greater peptide-peptide interactions to form either helix-rich (salt-free solutions) or β-sheet-rich (salt solutions) oligomers. Peptide-peptide and peptide-solvent interactions contribute to structural transitions, peptide self-assembly and oligomerization. These interactions were further investigated to understand their roles in aggregation kinetics of U3.5 peptide.

**Figure 6.**
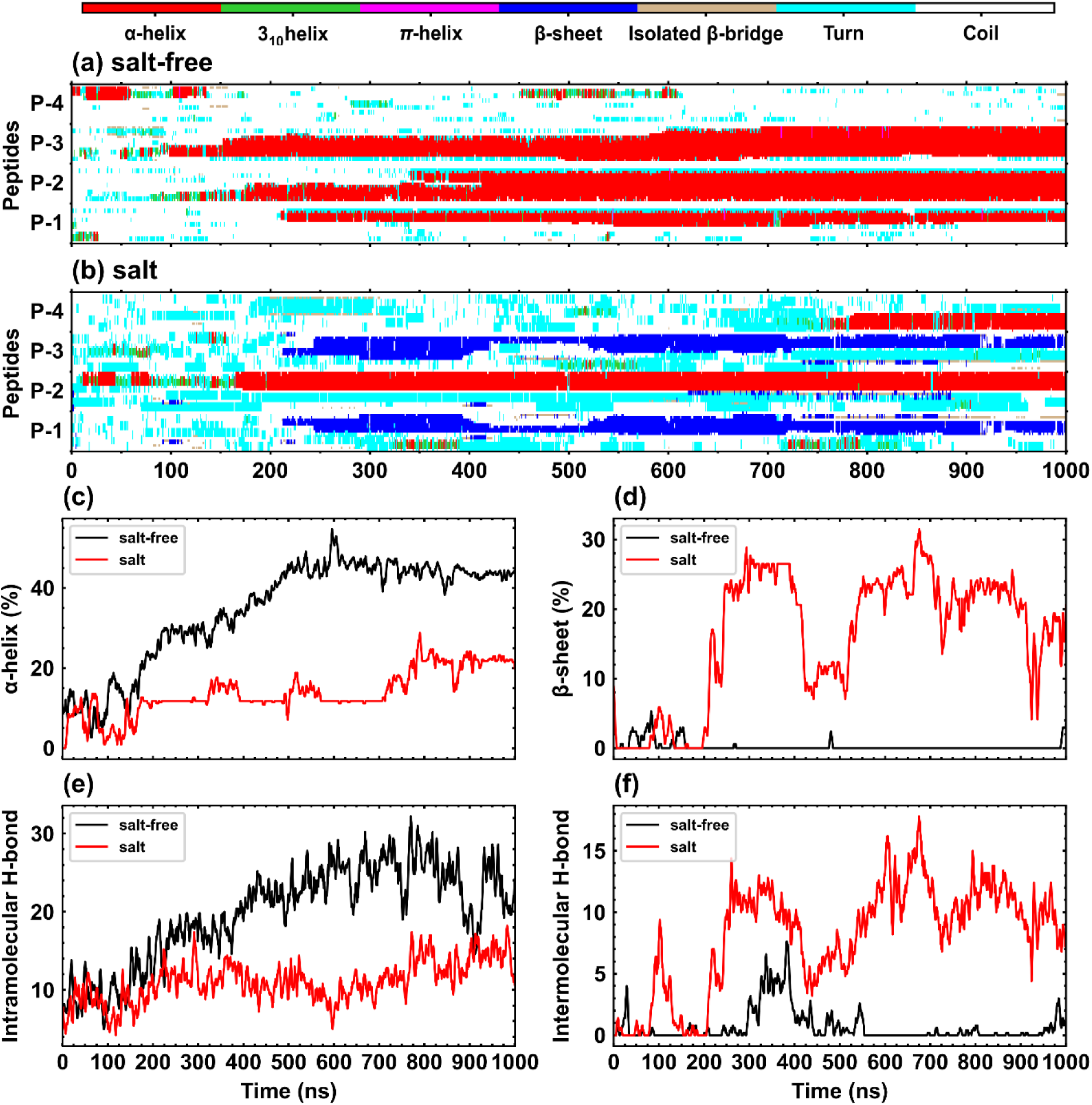
Structural transitions and the number of hydrogen bonds of helical intermediates formed in the tetramer simulations, in 10% TFE. A heat map showing the secondary structure status of peptides as they evolve into oligomers; (a) helical intermediates formed around 200 ns in salt-free condition showed the sustained presence of helices (widening of red strips) in oligomer evolution, (b) whereas the helical intermediate formed in presence of salt showed β-sheets transitions in P1 and P3 peptides. (c) and (d) Secondary structure transition over simulation trajectories of oligomers. The number of intra- and intermolecular hydrogen bonds of peptide oligomers in (e) and (f), respectively.

### Peptide-peptide interaction

#### Secondary structure and hydrogen bond

Secondary structure transitions and the number of hydrogen bonds, for simulations with 10% TFE, were quantified to understand the difference in oligomer evolution in the absence and presence of salts, respectively (Figure 6). In both cases, secondary structure transitions were similar until the helical intermediate began to form (Figures 6c, d at ≈ 200 ns). Once helical intermediates formed in these simulations, subsequent secondary structure transitions were qualitatively different in the salt-free and salt solutions. This was because salt modified peptide-peptide interactions, which in turn influenced both secondary structure transitions and peptide aggregation. In salt-free solutions, a sharp increase in the α-helix content of helical intermediates was observed, from ≈ 30% at 200 ns to nearly 50% at 600 ns. A very small increase in the α-helix content of the helical intermediate (from 15% at 200 ns to 20% at 1µs) was observed in presence of salt. However, a very large increase in β-sheet content was observed in the presence of salt (from nearly 0 to 25%). Intra- and inter-peptide interactions were investigated by quantifying the numbers of both intra- and inter-molecular H-bonds, respectively. The number of intra-molecular H-bonds induced during oligomerisation in salt-free media were almost double the number of intra-molecular H-bonds identified in oligomers formed in salt media. In contrast, the trend was reversed between the salt-free and salt solution for the inter-molecular H-bonds. A significant number of inter-molecular H-bonds formed in presence of salt corresponding to the 25% β-sheet content in the U3.5 oligomers for this case. However, peptide oligomers formed under salt-free conditions showed negligible inter-molecular H-bonds consistent with the absence of β-sheets in the absence of salt (Figure 6(f)). These results highlighted the differences in peptide-peptide interactions that ultimately led to formation of two different type of oligomers in salt-free and salt solutions, respectively.

#### Residue-residue interaction

The two types of oligomers, discussed above, showed opposite trends in intra- and inter-peptide interactions, indicating that residue-residue interactions were modified by the presence of salt. Interpeptide contact maps of clusters, at different stages of oligomer evolution, were generated to investigate residue-residue interactions. Figures 7(a) – (c) show contact maps for the 10% TFE salt-free case at different time instances during the simulation. Similarly, Figures 7(d) – (f) show contact maps for the 10% TFE case in presence of salt. In both cases, the contact maps (Figures 7(a) and (d)) showed strong residue-residue interactions toward the C-termini during helical intermediate formation. At later stages, when the oligomers evolved into compact structures (375 ns and 275 ns in salt-free and salt solutions, respectively); stronger residue-residue interactions were observed over the entire peptide. Whereas, residue-residue interactions were dominated by hydrophobic interactions in salt-free solutions (Figure 7(c)), both hydrophobic and hydrophilic interactions were significant in presence of salt in the solution (Figure 7(e)). Thus, the interpeptide contact maps showed a qualitative shift in residue-residue interactions at 10% TFE; from predominantly hydrophobic in salt-free solutions to a combination of hydrophilic and hydrophobic interactions in salt solutions. The contact maps in Figures 7(c) and (f) show that residue-residue interactions were also associated with the specific secondary structures that emerged in the oligomers. It is interesting to note that similar helical intermediates that formed in the early stages in the two cases (Figures 7 (a), (d)) evolved into α-helix (Figure 7 (c)) and β-sheet (Figure 7(f)) dominated oligomers in salt-free and salt solutions, respectively. Clearly, the change in residue-residue interactions upon addition of salt must be associated with a change in the solvent structure around the peptide clusters. This was determined by carefully examining peptide-solvent interactions both in absence and presence of salt.

**Figure 7.**
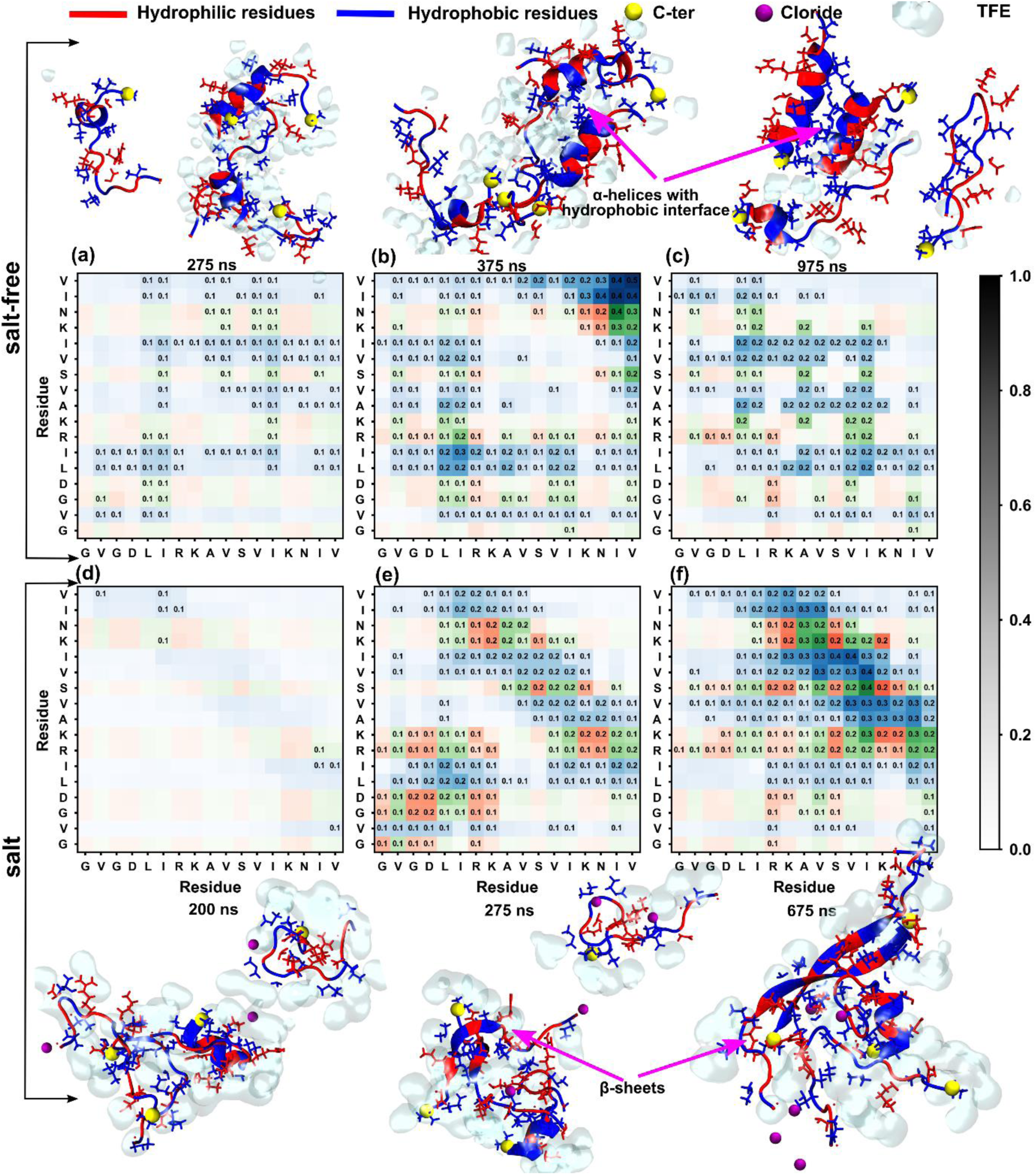
Interpeptide contact maps and corresponding snapshots of the helical intermediates evolved in 10% TFE, simulated in salt-free and salt solutions. Three stages of secondary structure evolution were taken to investigate residue-residue interactions, corresponding to structural transitions; initial, intermediate and later stages (a, b and c corresponding to 275, 375, and 975 ns for salt-free trajectory and d, e and f corresponding to 200, 275, and 675 ns for salt-containing simulation). In both the cases, residues toward the C-terminus interacted strongly to initiate the helical intermediate formation (a, d). During compact structure formation, acquired for the helical intermediates, the residue-residue interactions distributed throughout the peptides; predominant hydrophobic interaction in salt-free simulation (b), whereas both hydrophobic and hydrophilic interactions in salt-containing simulations (d), which similarly remained in later stages of helical intermediates (c and f). Interestingly, the evolution of helical intermediate in salt-free simulations were mainly driven by hydrophobic interactions whereas, both the hydrophobic and hydrophilic interactions prevailed in salt-containing simulations.

#### Peptide-solvent interactions

The specific association of TFE and water with different residues of U3.5 modulates inter-peptide interactions and determines structural evolution of clusters in TFE:water mixtures. Solvent-residue interactions were examined in detail for 10% TFE (both in salt-free and salt solutions) where fibrillation kinetics was faster compared to that in buffer. It was important to examine solvent-residue interactions at other TFE concentrations too. This was because peptide-solvent interactions changed with TFE concentration and influenced both secondary structure populations (Figure 4) and U3.5 aggregation propensity (Figure 5).

Radial distribution functions (RDFs) were evaluated to examine solvent distribution around the peptides. In the monomer simulations, TFE and water associated preferably with hydrophobic and hydrophilic residues, respectively (Figure 3). RDF plots of fluorine atom distribution (belonging to TFE) with respect to every hydrophobic residue (Figure 8) were generated for salt-free and salt solutions, respectively, at 400 ns in 10% TFE. In the salt-free case, RDF plots showed a stronger association of TFE molecules with hydrophobic residues towards N-termini (peak height between 2.75 – 3.25, Figure 8(a)) compared to residues towards the C-termini (peak height between 2.0 – 2.5, Figure 8(c)). In contrast, a reverse trend in residue-TFE interactions was observed in the presence of salt. The hydrophobic residues towards C-termini showed stronger association with TFE (peak height between 2.75 – 3.0, Figure 8(f)) compared to N-termini residues (peak height between 1.75 – 2.5, Figure 8(d)). A major difference in RDF can be seen between the salt-free and salt solutions with respect to the N-termini residues, where the peak height reduced from 3 for the salt-free case to 2 for the salt solution. These changes in RDF indicated the crucial role of salt in determining TFE association with hydrophobic residues, and by extension its role in modifying peptide-peptide interactions.

**Figure 8.**
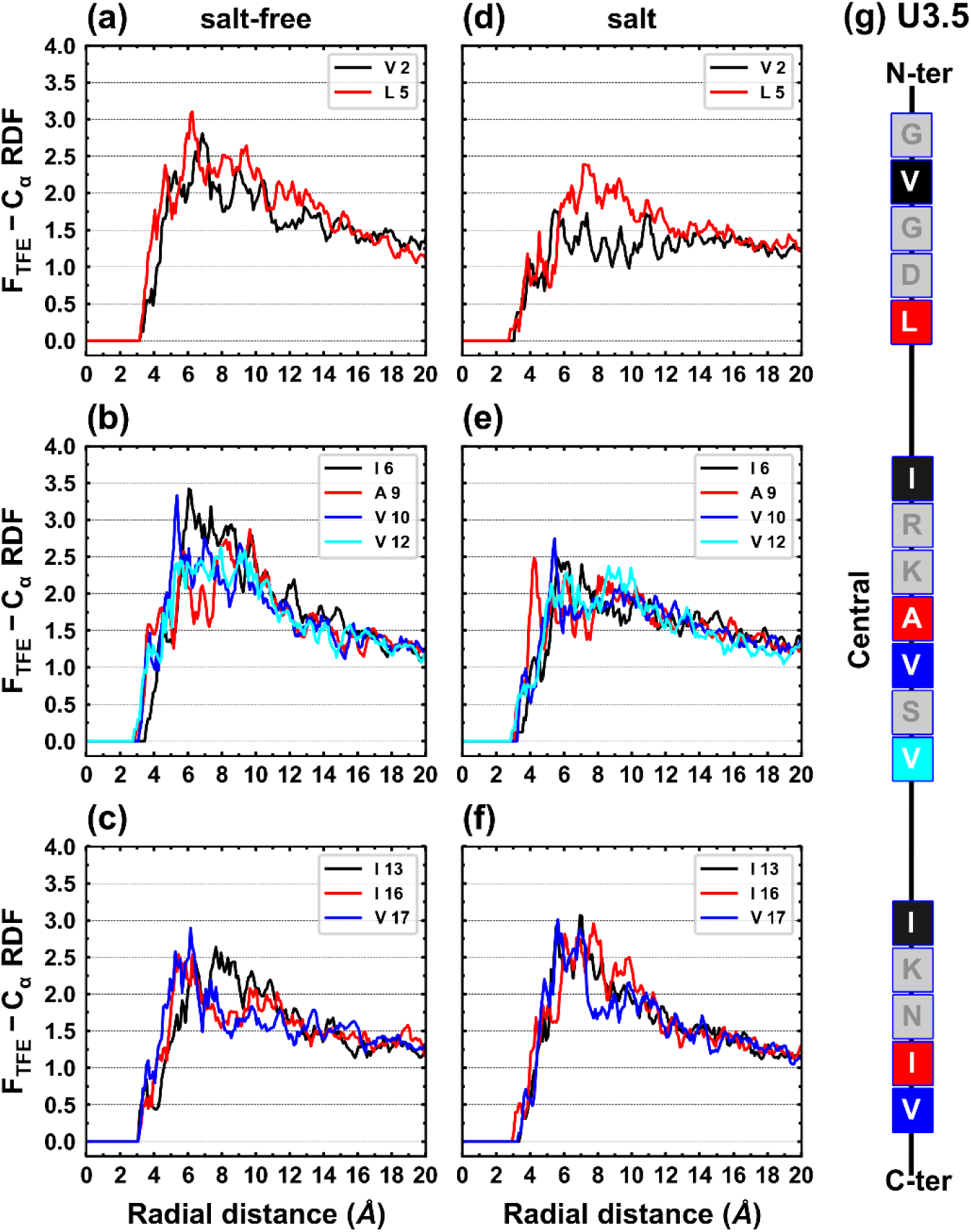
Radial distribution function (RDF) of individual hydrophobic residues of U3.5 of 10% TFE multi-peptide simulations in salt-free and salt solutions. For convenience, the U3.5 is illustrated into three segments; N-termini, central and C-termini in 10% TFE. The hydrophobic residues toward N-terminus show a strong association with TFE in salt-free media (peak height between 2.75 – 3.25) compared to salt media (peak height between 1.75 – 2.75). Significant differences also appear in RDF of central and C-termini segments of U3.5 compared to slat-free and salt media simulations.

The RDFs corresponding to the distribution of salt (chloride ions) around charged residues indicated strong interactions of ions with positively charged residues (Figures 9(a) and (b)). Since, most of the charged residues of U3.5 peptides are located near the N-terminus, therefore the effect of ions on TFE association with the peptide residues was more prominent near the N-termini. The presence of ions displaced the TFE molecules (Figure 8 (d)) closer to the N-termini. These results clearly indicated that one of the major roles of salt was to alter the TFE and water structure around peptides in a cluster. The impact of salt in reducing TFE-hydrophobic residue interactions was crucial for the secondary structure transitions in peptide clusters. Next, the accessible surface area (ASA), which quantified the exposed surface area of the peptide as a function of solvent composition, was calculated. The ASA values were averaged over the respective salt-free and salt solution trajectories (at 10% TFE) to compare the overall TFE-water interactions (Figure 9 (c) and (d)).

**Figure 9.**
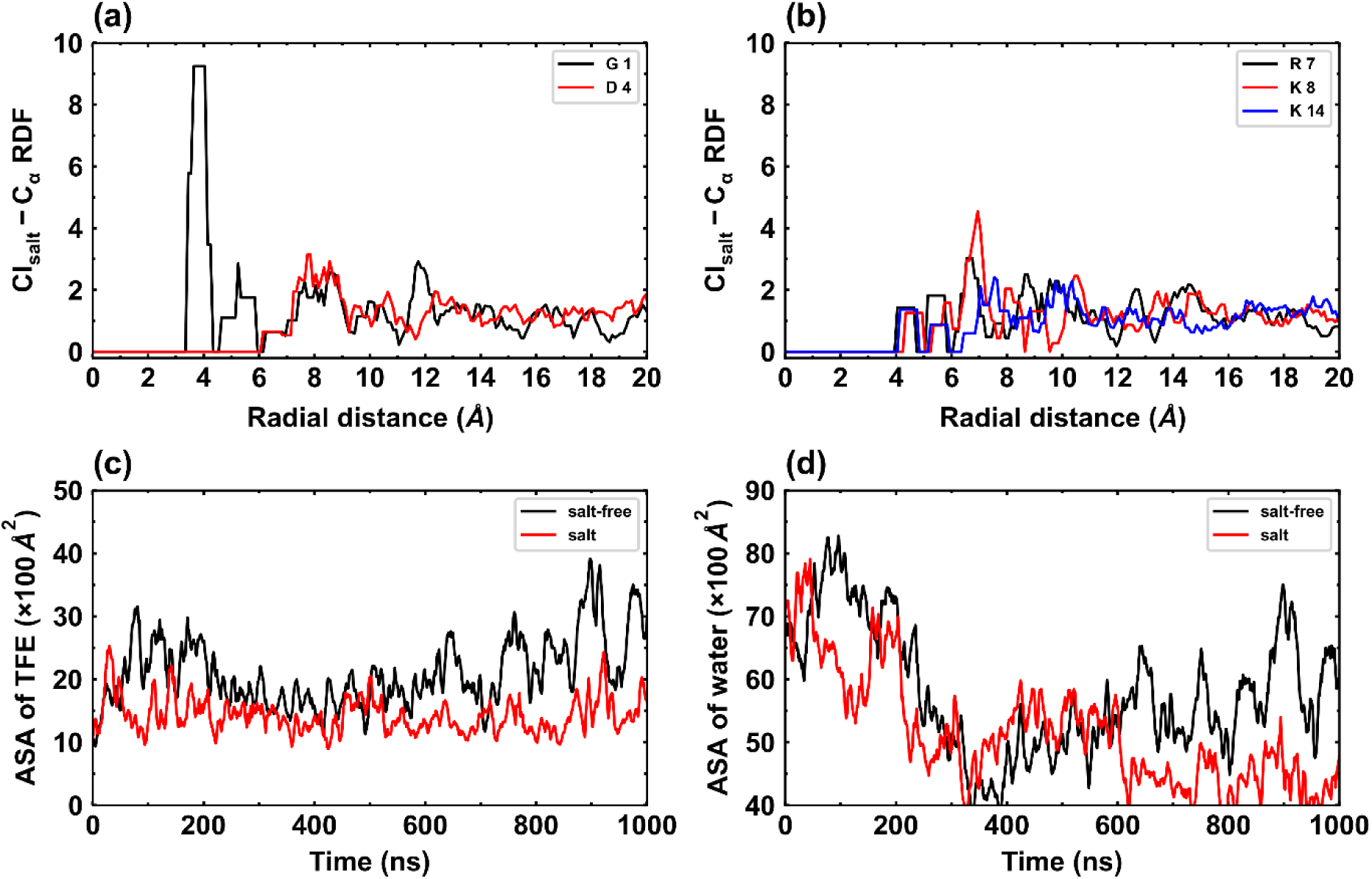
Radial distribution function (RDF) of chloride ions around individual charged residues (a and b) and accessible surface area (ASA) of peptide to solvents; TFE (c) and water (d). Chloride ions are strongly associated with positively charged residues, especially which are situated towards N-terminal. The ASA of TFE is significantly high in salt-free compared to salt, indicated the strong TFE interaction between peptide and TFE in salt-free simulations. The ASA of water showed similar trend over 600 ns of trajectories, however at the later-stage, the ASA of salt-free simulation is increased due to disintegration of its helical intermediate.

ASA values for both TFE and water were consistently lower in salt solutions when compared to the salt-free case. Chloride ions associated strongly with positively charged residues and displaced the TFE (or water) near the peptides compared to those in salt-free conditions. Furthermore, salt also plays a role in charge screening leading to a reduction in the electrostatic interactions within the peptide cluster.^42,43^ Our previous research on U3.5 peptide also revealed the role of salt in screening of charges.^44^ Charge screening effects can be inferred from the differences in residue-residue interactions in the contact maps in Figure 7. The contact maps, corresponding to salt solutions, showed enhanced residue-residue interaction near the N-terminus. Unlike in salt-free solution, positively charged residues at the N-termini showed greater interaction due to charge screening between like-charged residues. In 10% TFE, β-sheet transitions observed in peptide cluster primarily resulted from strong peptide-peptide interactions, which was facilitated by charge screening in presence of salt.

### Conclusions

Many intrinsically disordered peptides show the appearance of transient helical structures during the formation of β-sheet-rich amyloid fibrils. Several membrane mimetics, including TFE, modulate the secondary structures and aggregation kinetics of amyloid forming peptides.^15,28,45,46^ However, the molecular mechanism of this modulation is still not clear. In the current study, the role of TFE in determining structural transitions and aggregation kinetics of U3.5 in TFE:water mixtures was investigated using both biophysical experiments and molecular dynamics simulations. Further, the important role of the salt was also examined by studying peptide aggregation both in salt-free and salt-containing TFE:water mixtures. The fluorescence ThT assay results (Figure 1) showed that U3.5 peptide aggregation behaviour was dependent on the percentage of TFE present in water. At low TFE concentrations, aggregation of U3.5 increased 5 – 7 times when compared to the aggregation observed in saline buffer (no TFE present). However, this increase in aggregation did not begin until the addition of saline buffer. Furthermore, the CD experiments, conducted under similar conditions to the fluorescence assay, indicated the crucial role of helical structural transition prior to peptide self-assembly, leading to amyloid formation. In Figure 1, the equilibrium state observed in the aggregation kinetics was achieved at around 10 hrs after buffer addition, which indicated a progression of ‘invisible’ peptide clusters that assembled into amyloid.^36,47^ However, the CD results showed that the transition of U3.5 into β-sheet-rich structures was rapid. In Figure 2(b), U3.5 transitioned from a partial helical structure into β-sheet structures within 10 minutes. However, the process of self-assembly continued for another 10 hrs until an equilibrium state was achieved. Furthermore, the addition of 40 % TFE immediately led to disintegration of the amyloid fibrils yet, the transition of β-sheet structures to helical structures took 23hrs. These delays in structural transitions and of aggregation/disintegration processes could be the result of oligomer populations which were formed quickly and progressed rapidly on the self-assembly pathway. To understand at a molecular level, MD simulations of the U3.5 peptide were carried out using different TFE percentages (0 – 40% v/v) in salt and salt-free conditions. These simulations, Figures 3 and 4, showed an increased tendency of α-helix formation in U3.5 peptide with increasing percentage of TFE present in water. Whereas, the hydrophobic residues of the U3.5 peptide preferentially interacted with TFE molecules, the hydrophilic residues interacted with water molecules. Interestingly, the greatest extents of U3.5 aggregation into higher order clusters (Figure 5) were observed for the 10% TFE simulations, both in salt-free and salt solutions. These simulation results were consistent with the fluorescence experiments (Figure 1) where the maximum aggregation and fastest kinetics were also observed for the 10% TFE. For 10% TFE simulations in salt-free media, peptides formed clusters with helices as the dominant secondary structure. However, in presence of salt in the 10% TFE simulations, U3.5 clusters with predominantly β-sheets were observed. A detailed investigation of the structural evolution of different clusters, formed in salt and salt-free simulations, revealed the differential association of TFE at the N-termini of the U3.5 peptide (Figure 8). In salt-containing solutions, chloride ions interacted strongly with the positively charged residues located toward the N-termini of the peptide (Figure 9) and screened like-charge repulsions between different peptides. This increased residue-residue interactions towards the N-termini (Figure 7) of peptides and facilitated subsequent peptide-peptide interactions within a cluster. Enhanced inter-peptide interactions resulted in stronger aggregation and enabled conformational transitions into β-sheet-rich structures in U3.5 peptide (Figure 6).

U3.5 is an antimicrobial peptide which is also amyloidogenic. Like most AMPs, it adopts helical conformations when embedded in lipid membranes or micelles. TFE is a known α-helical stabilizer and a membrane mimetic. Hence, it is expected to stabilize helical conformations in U3.5. U3.5 is also amyloidogenic and membrane-peptide interactions are widely implicated in amyloid formation. In the current study, U3.5 displayed a range of aggregation behaviours and qualitatively different structural transitions as a function of TFE concentration in TFE:water mixtures. Experiments and MD simulations showed that enhanced U3.5 aggregation kinetics at low TFE concentrations (5 – 10% TFE) resulted from more stable helical intermediates (due to TFE) and greater inter-peptide interactions. The presence of salt in the solution further enhanced peptide-peptide interactions through electrostatic screening and contributed to conformational transition to β-sheet dominated clusters. In the context of peptide-membrane interactions, TFE:water mixtures highlighted the role of hydrophobic environments in stabilizing early-stage helical intermediates that arise in the pathway to formation of β-sheet rich amyloids.

## Acknowledgements

The authors acknowledge financial support from the Department of Biotechnology (DBT), Government of India (GOI) for project (IMURA0781) at IITB-Monash research academy and for the award of a PhD stipend to AKP.

## References

1. Chiti F, Dobson CM. Protein misfolding, amyloid formation, and human disease: a summary of progress over the last decade. Annu Rev Biochem. 2017;86:27–68.

2. Selkoe DJ. The molecular pathology of Alzheimer’s disease. Neuron. 1991;6(4):487–498.

3. Morris AM, Watzky MA, Finke RG. Protein aggregation kinetics, mechanism, and curve-fitting: A review of the literature. Biochim Biophys Acta - Proteins Proteomics. 2009;1794(3):375–397. 10.1016/j.bbapap.2008.10.016

4. Bhak G-B, Choe Y-J, Paik S-R. Mechanism of amyloidogenesis: nucleation-dependent fibrillation versus double-concerted fibrillation. BMB Rep. 2009;42(9):541–551.

5. Xi W-H, Wei G-H. Amyloid-β peptide aggregation and the influence of carbon nanoparticles*. Chinese Phys B. 2016;25(1):18704. doi:10.1088/1674-1056/25/1/018704

6. Sengupta U, Nilson AN, Kayed R. The Role of Amyloid-β Oligomers in Toxicity, Propagation, and Immunotherapy. EBioMedicine. 2016;6:42–49. doi:10.1016/j.ebiom.2016.03.035

7. Cline EN, Bicca MA, Viola KL, Klein WL. The Amyloid-β Oligomer Hypothesis: Beginning of the Third Decade. J Alzheimer’s Dis. 2018;64(s1):S567–S610. doi:10.3233/JAD-179941

8. Bemporad F, Chiti F. Protein misfolded oligomers: experimental approaches, mechanism of formation, and structure-toxicity relationships. Chem Biol. 2012;19(3):315–327.

9. Campioni S, Mannini B, Zampagni M, et al. A causative link between the structure of aberrant protein oligomers and their toxicity. Nat Chem Biol. 2010;6(2):140–147.

10. Cremades N, Cohen SIA, Deas E, et al. Direct observation of the interconversion of normal and toxic forms of α-synuclein. Cell. 2012;149(5):1048–1059.

11. Ladiwala ARA, Litt J, Kane RS, et al. Conformational differences between two amyloid β oligomers of similar size and dissimilar toxicity. J Biol Chem. 2012;287(29):24765–24773.

12. Mannini B, Mulvihill E, Sgromo C, et al. Toxicity of protein oligomers is rationalized by a function combining size and surface hydrophobicity. ACS Chem Biol. 2014;9(10):2309–2317. doi:10.1021/cb500505m

13. Vincenzi M, Mercurio FA, Leone M. About TFE: Old and New Findings. Curr Protein Pept Sci. 2019;20(5):425–451. doi:10.2174/1389203720666190214152439

14. Imai T, Kovalenko A, Hirata F, Kidera A. Molecular thermodynamics of trifluoroethanol-induced helix formation: Analysis of the solvation structure and free energy by the 3D-RISM theory. Interdiscip Sci Comput Life Sci. 2009;1:156–160.

15. Dammers C, Gremer L, Reiß K, et al. Structural analysis and aggregation propensity of pyroglutamate Aβ(3-40) in aqueous trifluoroethanol. PLoS One. 2015;10(11):1–11. doi:10.1371/journal.pone.0143647

16. Buck M. Trifluoroethanol and colleagues: cosolvents come of age. Recent studies with peptides and proteins. Q Rev Biophys. 1998;31(3):297–355.

17. Roccatano D, Colombo G, Fioroni M, Mark AE. Mechanism by which 2, 2, 2-trifluoroethanol/water mixtures stabilize secondary-structure formation in peptides: a molecular dynamics study. Proc Natl Acad Sci. 2002;99(19):12179–12184.

18. Walgers R, Lee TC, Cammers-Goodwin A. An indirect chaotropic mechanism for the stabilization of helix conformation of peptides in aqueous trifluoroethanol and hexafluoro-2-propanol. J Am Chem Soc. 1998;120(20):5073–5079.

19. Culik RM, Abaskharon RM, Pazos IM, Gai F. Experimental validation of the role of trifluoroethanol as a nanocrowder. J Phys Chem B. 2014;118(39):11455–11461. doi:10.1021/jp508056w

20. Muller I, Sarramégna V, Milon A, Talmont FJ. The N-terminal end truncated mu-opioid receptor: from expression to circular dichroism analysis. Appl Biochem Biotechnol. 2010;160:2175–2186.

21. Haney EF, Vogel HJ. NMR of antimicrobial peptides. Annu reports NMR Spectrosc. 2009;65:1–51.

22. Bradford AM, Bowie JH, Tyler MJ, Wallace JC. New antibiotic uperin peptides from the dorsal glands of the Australian Toadlet Uperoleia mjobergii. Aust J Chem. 1996;49(12):1325–1331. doi:10.1071/CH9961325

23. Baltutis V, O’Leary PD, Martin LL. Self-Assembly of Linear, Natural Antimicrobial Peptides: An Evolutionary Perspective. Chempluschem. 2022:e202200240.

24. John T, Dealey TJA, Gray NP, et al. The Kinetics of Amyloid Fibrillar Aggregation of Uperin 3.5 Is Directed by the Peptide’s Secondary Structure. Biochemistry. 2019;58(35):3656–3668. doi:10.1021/acs.biochem.9b00536

25. Calabrese AN, Liu Y, Wang T, et al. The Amyloid Fibril-Forming Properties of the Amphibian Antimicrobial Peptide Uperin3.5. ChemBioChem. 2016;17(3):239–246. doi:10.1002/cbic.201500518

26. Prasad AK, Tiwari C, Ray S, et al. Secondary Structure Transitions for a Family of Amyloidogenic, Antimicrobial Uperin 3 Peptides in Contact with Sodium Dodecyl Sulfate. Chempluschem. 2022;87(1):e202100408.

27. Prasad AK, Tiwari C, Ray S, et al. Secondary Structure Transitions for a Family of Amyloidogenic, Antimicrobial Uperin 3 Peptides in Contact with Sodium Dodecyl Sulfate. Chempluschem. 2022;87(1). doi:10.1002/cplu.202100408

28. Calabrese AN, Liu Y, Wang T, et al. The Amyloid Fibril-Forming Properties of the Amphibian Antimicrobial Peptide Uperin3.5. ChemBioChem. 2016;17(3):239–246. doi:10.1002/cbic.201500518

29. Chiti F, Dobson CM. Protein Misfolding, Functional Amyloid, and Human Disease. Annu Rev Biochem. 2006;75(1):333–366. doi:10.1146/annurev.biochem.75.101304.123901

30. Stefani M. Protein misfolding and aggregation: new examples in medicine and biology of the dark side of the protein world. Biochim Biophys Acta (BBA)-Molecular Basis Dis. 2004;1739(1):5–25.

31. Kirkitadze MD, Condron MM, Teplow DB. Identification and characterization of key kinetic intermediates in amyloid $β$-protein fibrillogenesis. J Mol Biol. 2001;312(5):1103–1119.

32. Fezoui Y, Teplow DB. Kinetic studies of amyloid $β$-protein fibril assembly: differential effects of $α$-helix stabilization. J Biol Chem. 2002;277(40):36948–36954.

33. Kim B, Do TD, Hayden EY, Teplow DB, Bowers MT, Shea JE. Aggregation of chameleon peptides: Implications of α-helicity in fibril formation. J Phys Chem B. 2016;120(26):5874–5883. doi:10.1021/acs.jpcb.6b00830

34. Williamson JA, Miranker AD. Direct detection of transient $α$-helical states in islet amyloid polypeptide. Protein Sci. 2007;16(1):110–117.

35. Housmans JAJ, Wu G, Schymkowitz J, Rousseau F. A guide to studying protein aggregation. FEBS J. 2023;290(3):554–583.

36. Prasad AK, Martin LL, Panwar AS. Helical intermediate formation and its role in amyloids of an amphibian antimicrobial peptide. Phys Chem Chem Phys. 2023;25(17):12134–12147. doi:10.1039/D3CP00104K

37. John T, Dealey TJA, Gray NP, et al. The kinetics of amyloid fibrillar aggregation of uperin 3.5 is directed by the peptide’s secondary structure. Biochemistry. 2019;58(35):3656–3668.

38. Xue C, Lin TY, Chang D, Guo Z. Thioflavin T as an amyloid dye: fibril quantification, optimal concentration and effect on aggregation. R Soc open Sci. 2017;4(1):160696.

39. Morales R, Moreno-Gonzalez I, Soto C. Cross-seeding of misfolded proteins: implications for etiology and pathogenesis of protein misfolding diseases. PLoS Pathog. 2013;9(9):e1003537.

40. John T, Gladytz A, Kubeil C, Martin LL, Risselada HJ, Abel B. Impact of nanoparticles on amyloid peptide and protein aggregation: a review with a focus on gold nanoparticles. Nanoscale. 2018;10(45):20894–20913.

41. Kim B, Do TD, Hayden EY, Teplow DB, Bowers MT, Shea J-E. Aggregation of Chameleon Peptides: Implications of α-Helicity in Fibril Formation. 2016. doi:10.1021/acs.jpcb.6b00830

42. Aramvash A, Seyedkarimi MS. All-atom molecular dynamics study of four RADA 16-I peptides: the effects of salts on cluster formation. J Clust Sci. 2015;26:631–643.

43. Hong Y, Pritzker MD, Legge RL, Chen P. Effect of NaCl and peptide concentration on the self-assembly of an ionic-complementary peptide EAK16-II. Colloids Surfaces B Biointerfaces. 2005;46(3):152–161.

44. Ray S, Holden S, Martin LL, Panwar AS. Mechanistic insight into the early stages of amyloid formation using an anuran peptide. Pept Sci. 2019;111(5):e24120.

45. Khan MS, Tabrez S, Bhat SA, Rabbani N, Al-Senaidy AM, Bano B. Effect of trifluoroethanol on α-crystallin: folding, aggregation, amyloid, and cytotoxicity analysis. J Mol Recognit. 2016;29(1):33–40.

46. John T, Piantavigna S, Dealey TJA, Abel B, Ris- HJ, Martin LL. Effects of Oxidized and Anionic Lipids on the Aggregation of Amy-loid Beta and Uperin 3 . 5 Peptide. :1–16.

47. Abedini A, Raleigh DP. A critical assessment of the role of helical intermediates in amyloid formation by natively unfolded proteins and polypeptides. Protein Eng Des Sel. 2009;22(8):453–459. doi:10.1093/protein/gzp036

