## Supplementary information for "The origin of secondary structure transitions and peptide self-assembly propensity in trifluoroethanol-water mixtures"

### Abbreviations

| | |
|---|---|
| SDS | Sodium dodecyl sulphate |
| POPC | Phosphatidylcholine |
| TFE | 2,2,2-TriFluoroEthanol |
| MD | Molecular Dynamics |
| ThT | Thioflavin T |
| Uperin 3.5 | U3.5 |
| CD | Circular dichroism |
| NMR | Nuclear Magnetic Resonance |
| PBS | Phosphate buffer saline |
| NaCl | Sodium chloride |
| RDF | Radial distribution function |
| $R_g$ | Radius of gyration |
| ASA | Accessible surface area |

**Supplementary Tables and Figures**

**Table S1.** Detailed information of simulation setups.

| | 0% TFE | 10% TFE | 15% TFE | 20% TFE | 40% TFE |
|---|---|---|---|---|---|
| **Monomer simulations (Without salt)** | | | | | |
| **Peptides No.** | 1 | | | | |
| **TFE atoms** | 0 | 173 | 263 | 355 | 750 |
| **Water molecules** | 6753 | 6228 | 5947 | 5680 | 4500 |
| **Total atoms** | 20533 | 20515 | 20482 | 20509 | 20524 |
| **Simulation time (ns)** | 500 | 500 | 500 | 500 | 500 |
| **Monomer simulations (Salt)** | | | | | |
| **Peptides No.** | 1 | | | | |
| **TFE atoms** | 0 | 173 | 263 | 355 | 750 |
| **Water atoms** | 6712 | 6191 | 5910 | 5645 | 4471 |
| **Total atoms** | 20451 | 20441 | 20408 | 20439 | 20466 |
| **Simulation time** | 500 | 500 | 500 | 500 | 500 |
| **Tetramer simulations (Without salt)** | | | | | |
| **Peptides No.** | 4 | | | | |
| **TFE atoms** | 0 | 816 | 1248 | 1675 | 3537 |
| **Water atoms** | 31532 | 29376 | 28090 | 26808 | 21224 |
| **Total atoms** | 95704 | 96568 | 96598 | 96595 | 96601 |
| **Simulation time** | 500 | 1000 | 500 | 500 | 500 |
| **Tetramer simulations (Salt)** | | | | | |
| **Peptides No.** | 4 | | | | |
| **TFE atoms** | 0 | 816 | N/A | 1675 | 3537 |
| **Water atoms** | 31339 | 29200 | N/A | 26646 | 21092 |
| **Total atoms** | 95301 | 96216 | N/A | 96271 | 96337 |
| **Simulation time** | 500 | 500 | N/A | 500 | 500 |

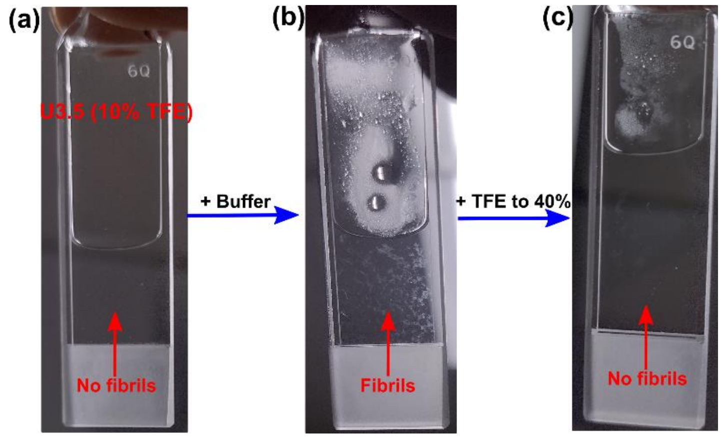


**Figure S1.** Images captured of the CD cuvette at different stages of experiments, that illustrate the deposited amyloid fibrils on the cuvette wall. (a) CD cuvette containing U3.5 peptide in 10% TFE-water, having no fibrils on wall (b) CD cuvette after 24 hrs of buffer addition, showing the smear of amyloid fibrils on walls (c) addition of TFE up to 40% led disappearance of amyloid fibrils from walls.

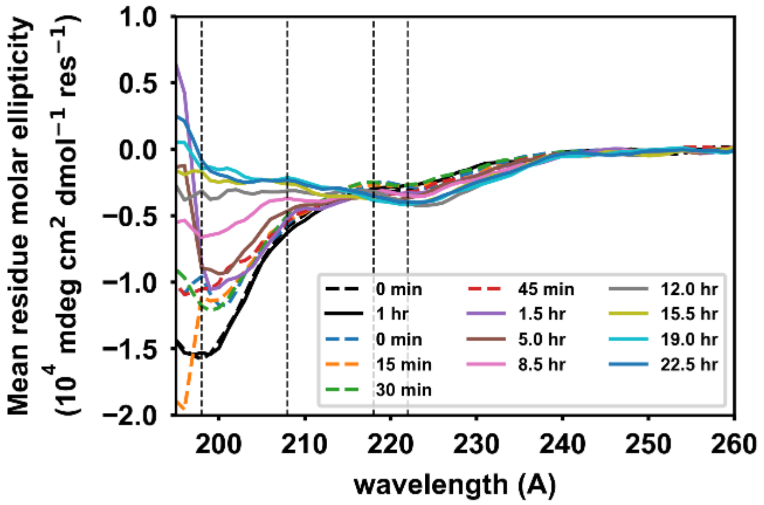


**Figure S2.** CD spectra of U3.5 peptide in PBS buffer. The time series spectra show the slow transitions of peptide secondary structure. In water, the U3.5 peptide remains in a random coil structure (black line). After the addition of buffer, the reduction of depth of the curve at 198 Å indicates the decrease of the random coil structure of the peptide. The spectra of 5 and 8.5 hrs show the mixed secondary structure (random coil, partial helical, and β-sheet), which means peptides transitioned into β-structure through the helical structure.

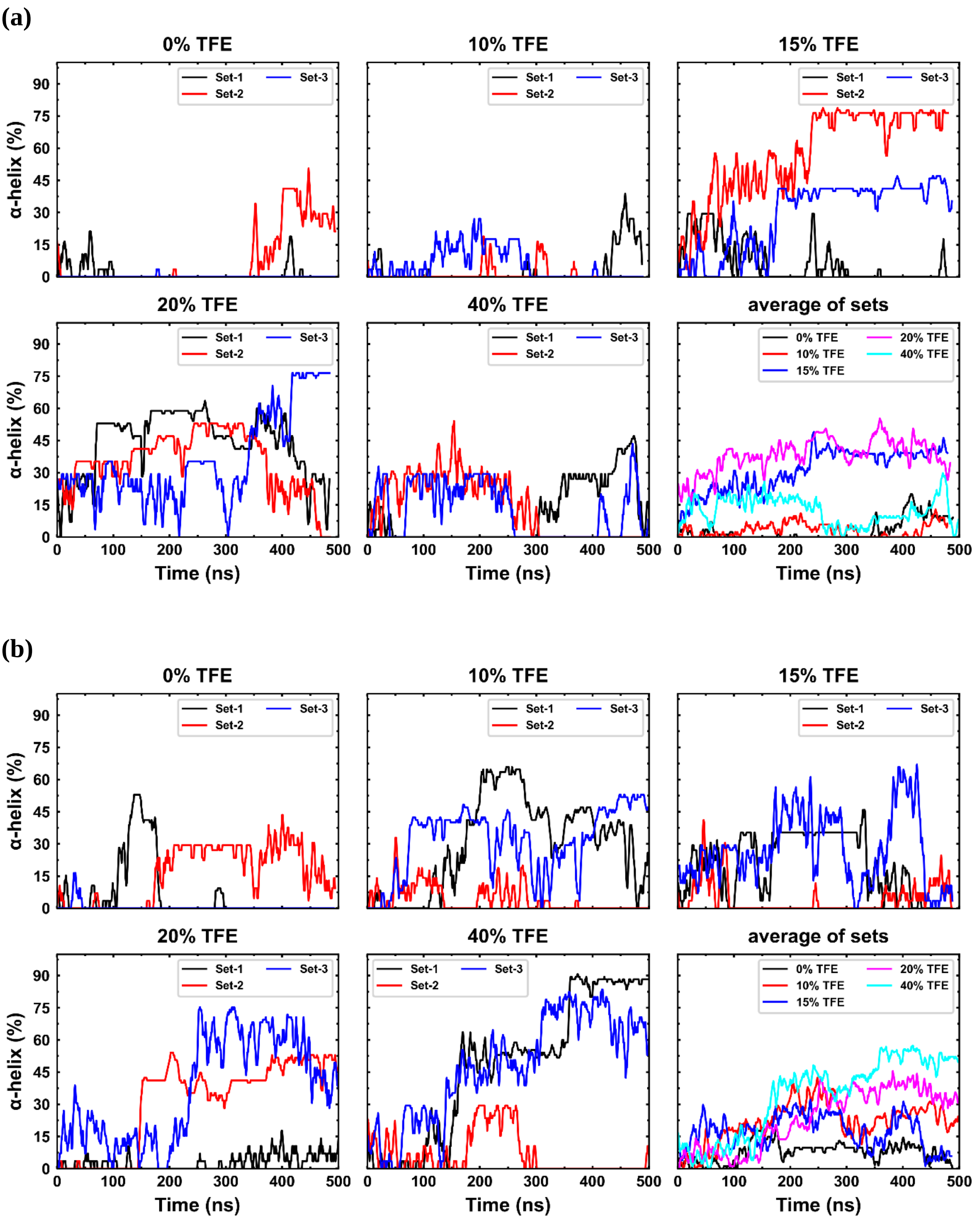


**Figure S3.** Evolutions of helical content in the U3.5 peptide of monomer simulations. Monomer simulations were run thrice, set-1, set-2 and set-3, in salt-free and salt media. The secondary structure of the peptide increased continuously over the trajectories in each set of simulations. The averages of sets show the increased helical content in peptide with incremental TFE percentage in solution (coherent in salt media).

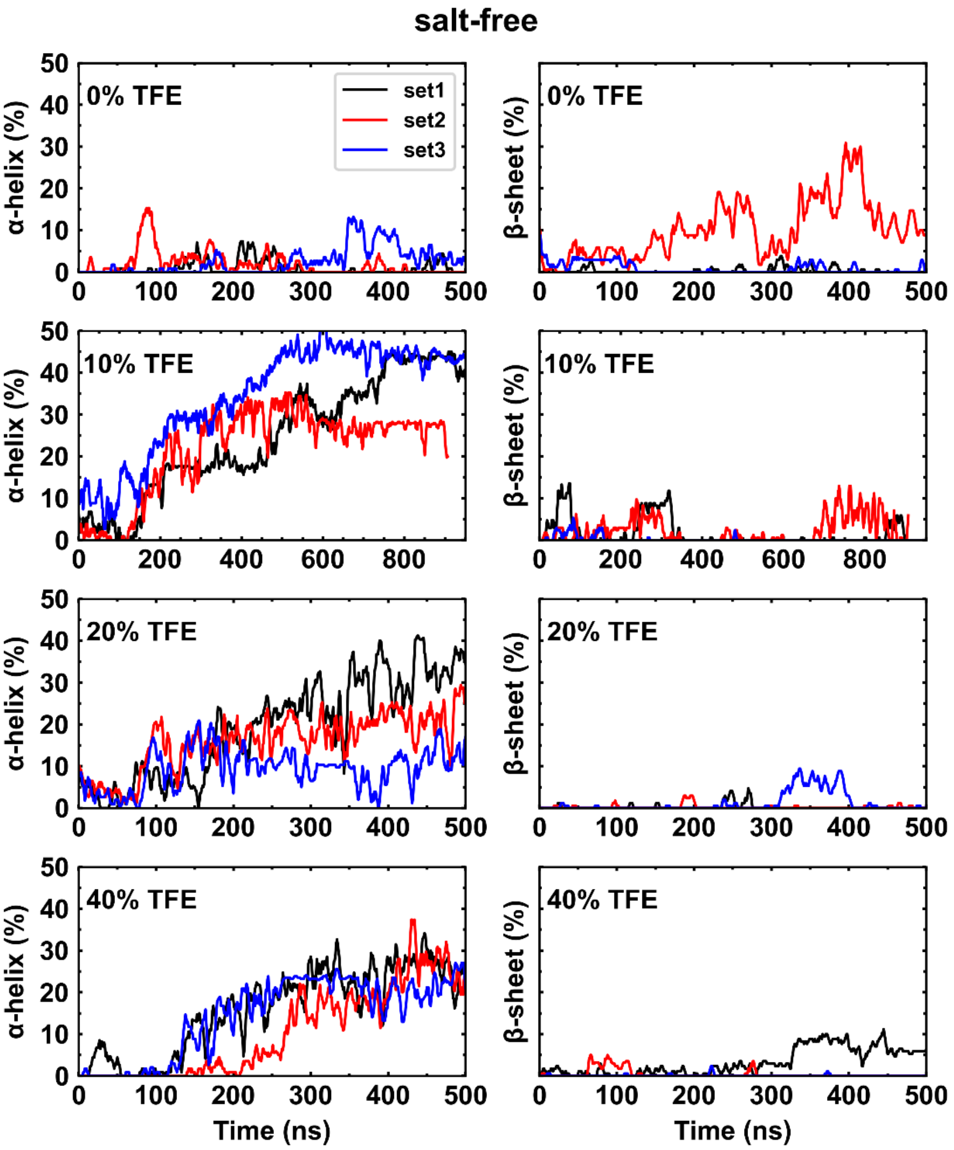
salt-free
0% TFE
10% TFE
20% TFE
40% TFE
set1
set2
set3
α-helix (%)
β-sheet (%)
Time (ns)

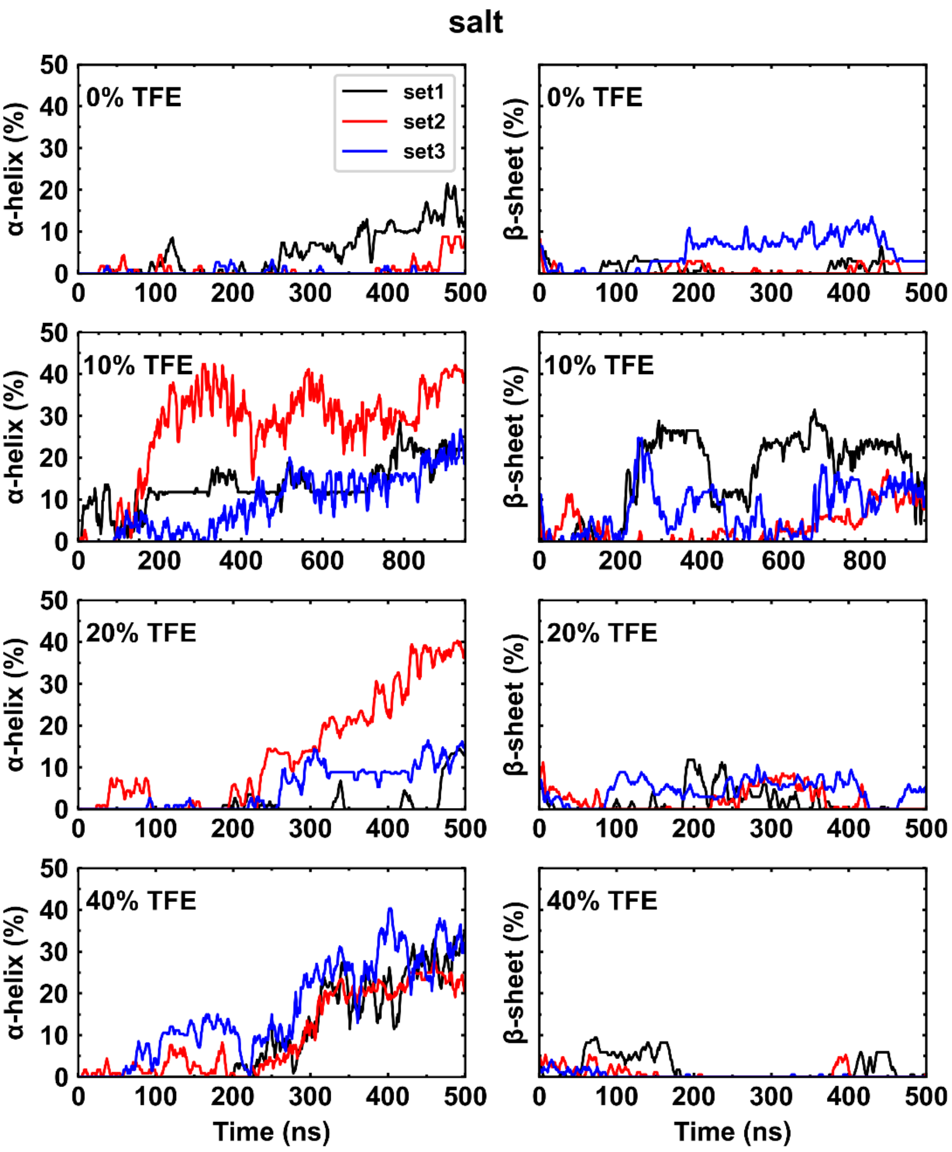


**Figure S4.** Evolutions of secondary structure in the U3.5 peptide of tetramer simulations at various TFE percentages. The percentage of α-helix content increased consistently in peptides both (a) salt-free and (b) salt media containing simulation. However, the β-sheet formation was mainly observed in simulations containing salt media, specifically at 10% TFE.

### salt-free

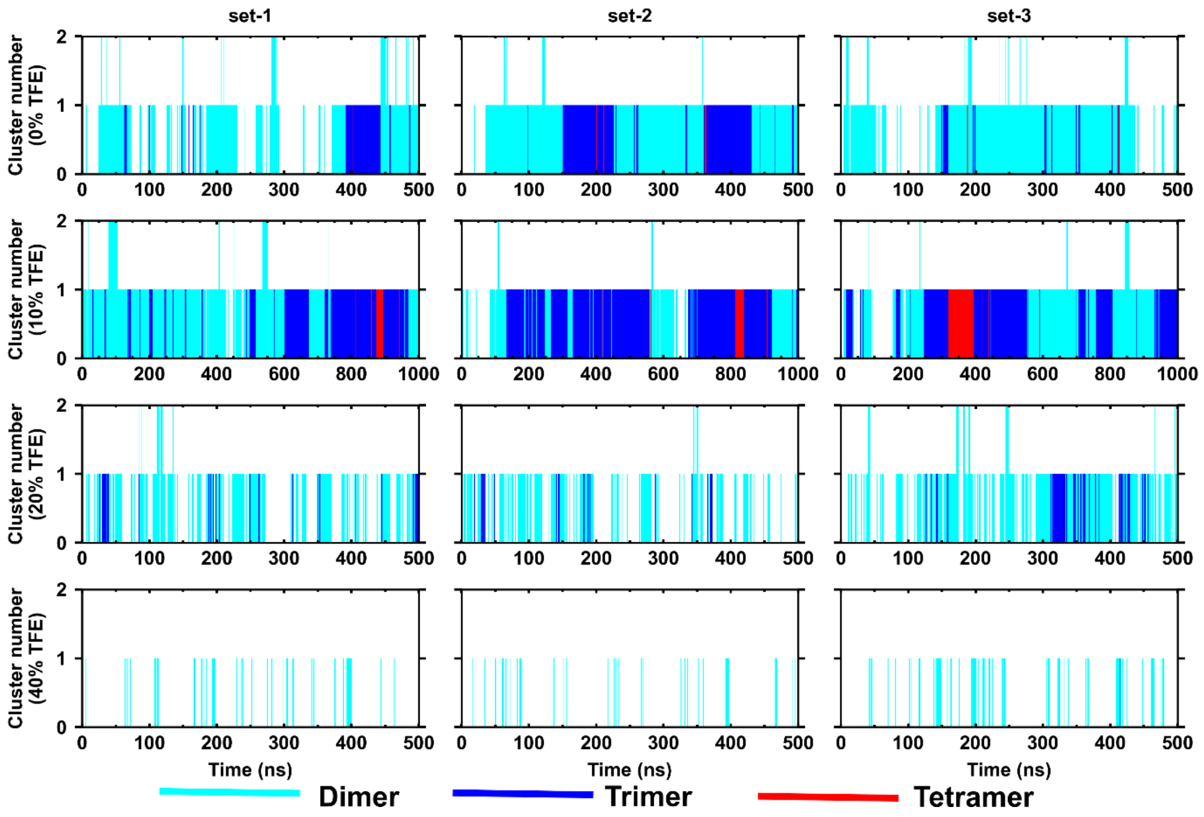


### salt

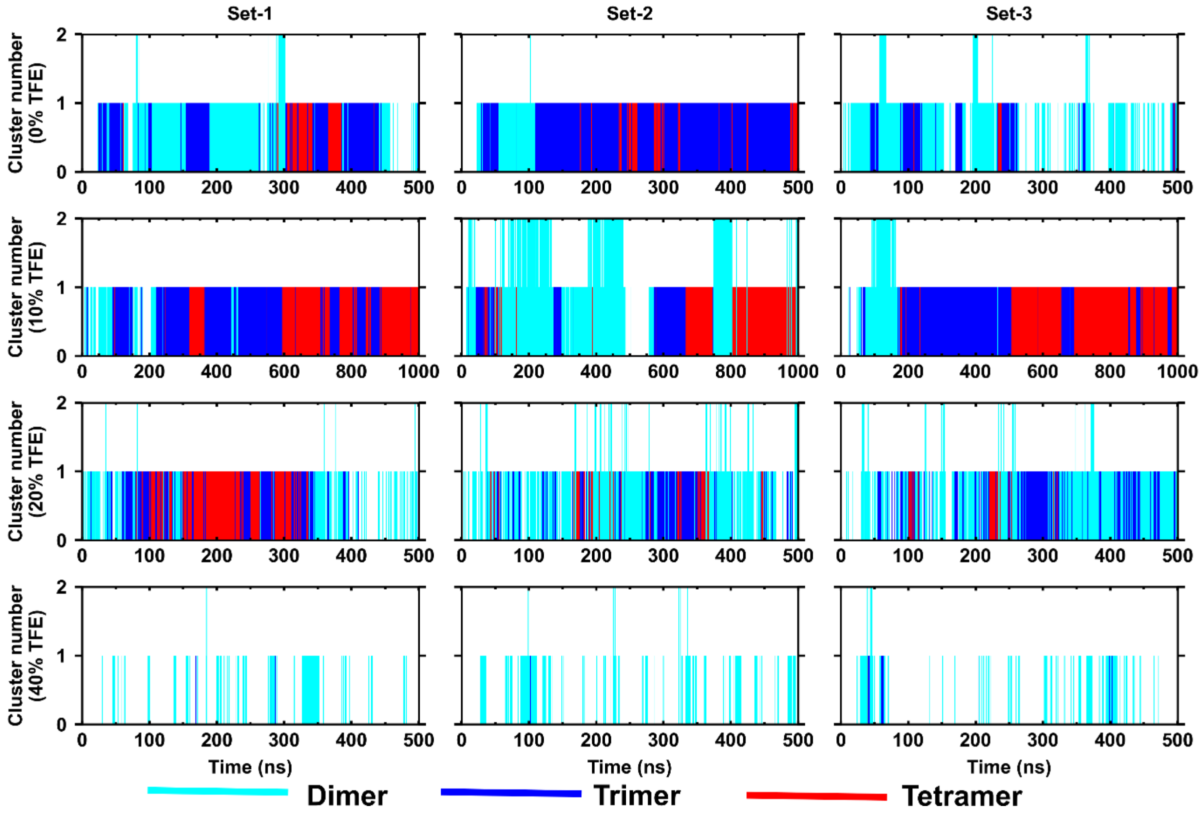

**Figure S5.** Peptide aggregation of tetramer simulations at various TFE percentages are shown here. The aggregations of peptides were analysed using cluster status, a number of peptides associated within a distance of 5Å. (a) In salt-free media, peptides formed the cluster mostly of trimeric structure in 10% TFE, whereas the other simulations, especially for 40% TFE content, showed reluctance to aggregations. (b) The salt containing media showed an increased tendency in peptide aggregations, highest at 10% TFE and complete inhibition at 40% TFE simulations.

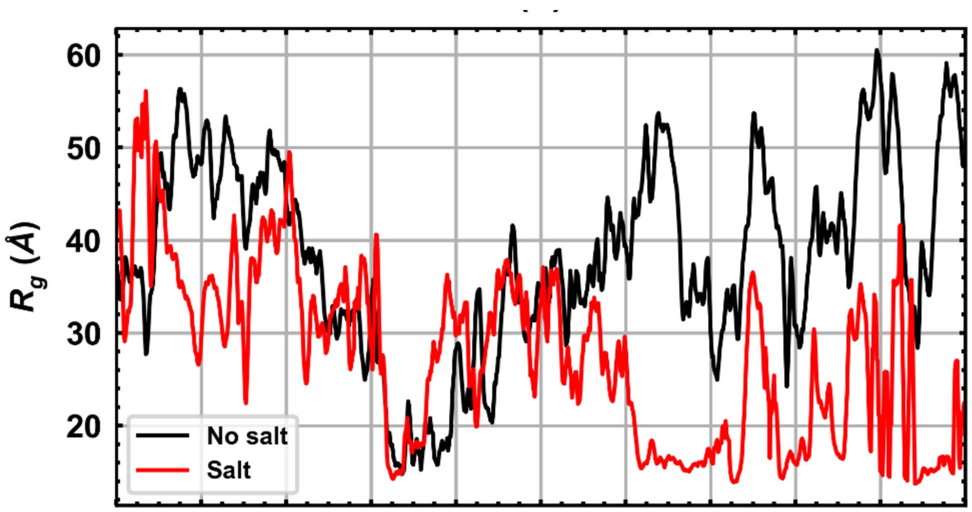


**Figure S6.** The radius of gyration of helical intermediates formed in 10% TFE simulations in salt-free and salt containing media.
